# An inactivated multivalent influenza A virus vaccine is broadly protective in mice and ferrets

**DOI:** 10.1101/2021.09.10.459807

**Authors:** Jaekeun Park, Sharon Fong, Louis M. Schwartzman, Zhong-Mei Sheng, Ashley Freeman, Lex Matthews, Yongli Xiao, Mitchell D. Ramuta, Natalia A. Batchenkova, Li Qi, Luz Angela Rosas, Stephanie Williams, Kelsey Scherler, Monica Gouzoulis, Ian Bellayr, David M. Morens, Kathie-Anne Walters, Matthew J. Memoli, John C. Kash, Jeffery K. Taubenberger

## Abstract

Influenza A viruses (IAVs) present major public health threats from annual seasonal epidemics, from pandemics caused by novel virus subtypes, and from viruses adapted to a variety of animals including poultry, pigs and horses. Vaccines that broadly protect against all such IAVs, so-called “universal” influenza vaccines, do not currently exist, but are urgently needed. This study demonstrates that an inactivated, multivalent whole virus vaccine, delivered intramuscularly or intranasally, is broadly protective against challenges with multiple IAV HA/NA subtypes in both mice and ferrets, including challenges with IAV subtypes not contained in the vaccine. This vaccine approach indicates the feasibility of eliciting broad “universal” IAV protection, and identifies a promising candidate for influenza vaccine clinical development.

**One-Sentence Summary:** An inactivated, whole avian influenza virus vaccine delivered intramuscularly or intranasally provides extremely broad protection against antigenically divergent viral challenge and is a promising candidate for a “universal” influenza virus vaccine.

## Introduction

Influenza A viruses (IAVs) pose a continual major public health threat. Globally, endemic (annual, or ‘seasonal’) influenza results in 3-5 million severe illnesses and up to 650,000 deaths each year (*1*). Influenza pandemics, in which novel IAVs unpredictably emerge from the IAV reservoir of wild waterfowl (*2*), and against which most humans lack protective immunity, can have even larger global impacts (*3*): e.g., the 1918 influenza pandemic resulted in at least 50 million deaths (*4*). In addition, IAVs adapted to non-human hosts emerge sporadically to infect and kill humans (e.g., poultry-associated H5N1 and H7N9) or even pandemically (e.g., pandemic H1N1 “swine” influenza in 2009). The fact that IAVs are permanently adapted to, or repeatedly infect, a wide variety of non-human hosts such as horses, dogs, seals, and other hosts indicates that IAV risks to humans are widely distributed in nature; moreover, these viruses are comprised of a broad array of different genotypes of variable and often unpredictable human pathogenicity. Currently, the only IAV vaccines licensed for human use are made each year to match specific circulating influenza virus strains in both Northern and Southern Hemispheres (*5*). Such vaccines do not protect against antigenically variant annual IAV strains, new pandemic viruses, poultry-associated viruses, or viruses adapted to, or frequently infecting, other mammalian hosts. There is a critical need for influenza vaccines that broadly protect against all such IAVs, a so-called “universal” vaccine (*6, 7*).

IAVs are enveloped, negative-sense, single-stranded RNA viruses with 8 genome segments (*8*). In addition to humans, IAVs infect large numbers of warm-blooded animal hosts, including over 100 avian species and many mammals (*9, 10*). IAVs express two major surface glycoproteins—HA and NA, and are subtyped by antigenic characterization of the HA and NA glycoproteins. Sixteen HA and 9 NA subtypes are consistently found in avian hosts in various combinations (e.g., A/H1N1 or A/H3N2), and these wild bird viruses are thought to be the ultimate source of human pandemic influenza viruses (*9*). IAV genome segmentation allows for viral reassortment, and since HA and NA are encoded on separate gene segments, novel IAVs of any of the 144 possible subtype combinations can theoretically be generated following mixed infections in a host, in a process called “antigenic shift”. IAVs are also evolutionarily dynamic RNA viruses with high mutation rates. Mutations that change amino acids in the antigenic portions of HA and NA proteins may allow human-adapted strains to evade population immunity, a process termed “antigenic drift”. Despite enhanced surveillance, future pandemics cannot yet be predicted, including when and where a pandemic virus strain will emerge, what the viral subtype will be, how pathogenic it will be in humans, or whether there will be some immunologic cross-reactivity with prior circulating IAVs. Severe human zoonotic infections with poultry-origin IAVs have also been observed, including recent human infections with A/H5N1 and A/H7N9 viruses (*11*).

An effective ‘universal’ vaccine would ideally provide broad protection against all IAV subtypes found in birds, domestic mammals, and humans. Efforts to develop such broadly protective vaccines have been under way for decades (*12*) and have included experimental vaccines specifically targeting the M2 ectodomain (*13, 14*) or NA (*15, 16*) proteins to stimulate the development of protective antibody responses, vaccines based on antigens that stimulate development of T-cell responses (*17*), and most recently, a variety of vaccine approaches targeting antigenically conserved epitopes on the HA head and stalk (*18–21*). However, a practical vaccine inducing broad heterosubtypic or universal protection has not been previously demonstrated with any of the above approaches.

In the present study, mice and ferrets vaccinated with a whole-virus beta-propiolactone (BPL) inactivated vaccine cocktail intranasally or intramuscularly were subsequently challenged with multiple IAVs. Challenge viruses included homosubtypic viruses (strains with antigenically variable HA and NA subtypes sharing vaccine virus subtypes, as examples of protection against annual influenza) and heterosubtypic viruses (strains with HA and/or NA subtypes not present in the vaccine, as examples of zoonotic and pandemic IAVs) viruses showed near 100% protection. The vaccine cocktail included four inactivated wild-type, low pathogenicity avian influenza viruses: H1N9, H3N8, H5N1, and H7N3. The four HA subtypes were chosen to reflect the subtypes of currently circulating annual IAV strains (H1 and H3) or recent epizootic IAV infections (H5 and H7) and represent both major phylogenetic HA groupings—clade 1 (H1 and H5) and clade 2 (H3 and H7) (*20*), along with four different NA subtypes representing the two major NA clades (*2*).

## Results

### Immune responses to IM and IN vaccination in mice and protection against homosubtypic and heterosubtypic viral challenge

The multivalent vaccine was prepared using BPL-inactivation of avian IAV H1N9, H3N8, H5N1, and H7N3 subtypes, grown in Madin-Darby Canine Kidney (MDCK) cells, and purified by sucrose density gradient. Mice were primed on day 0 and boosted 28 days later by intramuscular (IM) or intranasal (IN) vaccination with 6ug total protein (1.5ug per subtype). An overview of animal studies is depicted in Supplementary Fig. S1. The vaccine was highly immunogenic in mice and both IN and IM immunization elicited significant serum IgG antibody responses to homologous HAs (H1, H3, H5, H7) and NAs (N1, N3, N8, N9) as well as hemagglutination inhibition (HAI) antibodies (Fig. 1A,B, Supplementary Fig. S2A). Although IM immunization induced generally higher serum IgG antibody responses than IN immunization (Fig 1A,B), IN immunization induced a more pronounced IgA response in bronchoalveolar lavage (BAL) fluid (Supplementary Fig. S2B,C). Additionally, antibodies to the conserved stalk (or stem) regions of both group 1 and group 2 HAs were generated by IM or IN immunization (Supplementary Fig. S2D,E).

**Figure 1.**
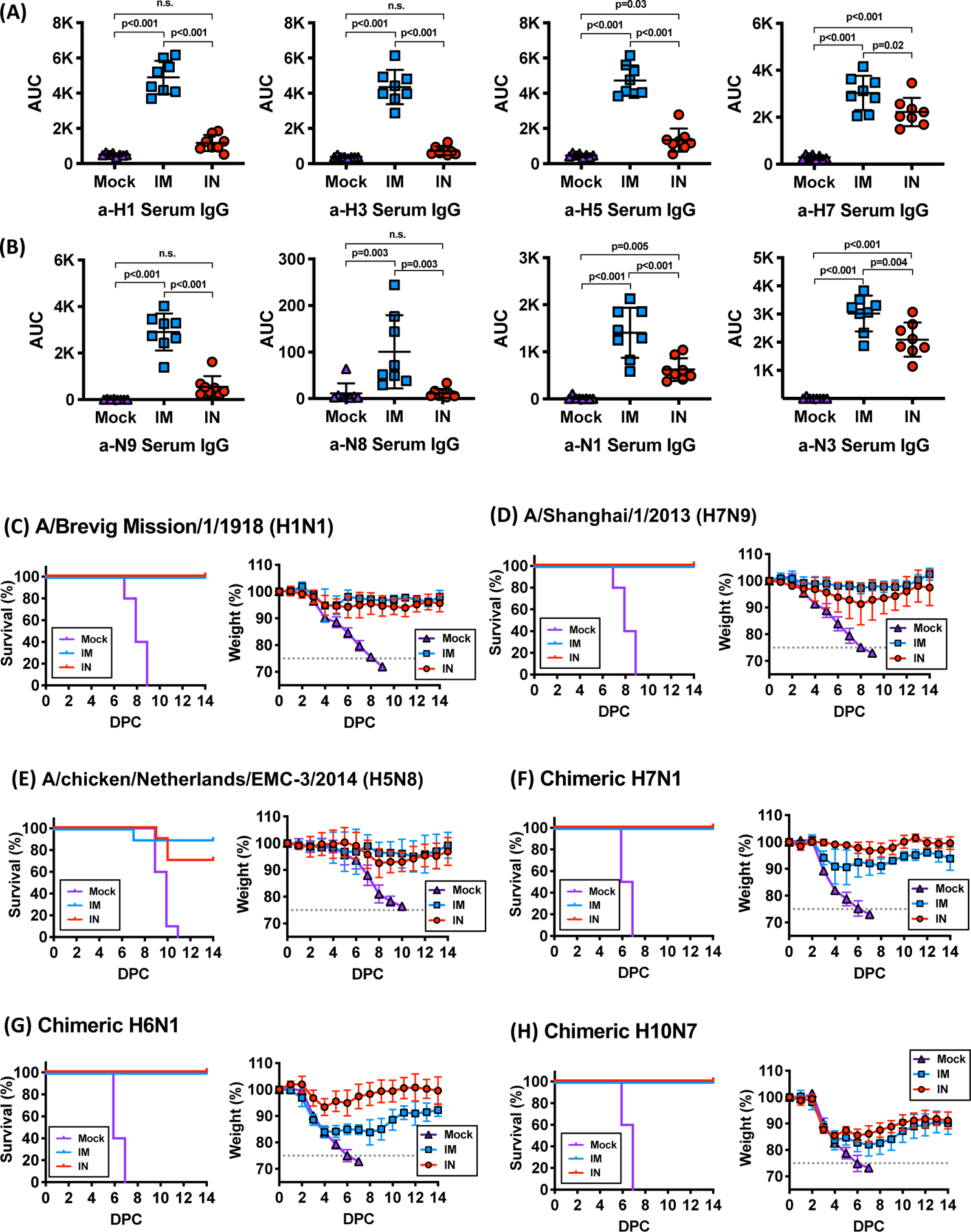
Immunogenicity and protective efficacy of the BPL-inactivated vaccine in mice. Serum IgG levels against (**A**) four vaccine hemagglutinin (HA) antigens and (**B**) four vaccine neuraminidase (NA) antigens were measured 3 weeks after the boost immunization in mock-, IM-, or IN-vaccinated mice using ELISA. Ordinary ANOVA test and post hoc Tukey’s multiple comparison test were used to compared antibody levels between groups. n.s; not significant **(C-F).** Percent survival and percent weight loss in mock-, IM-, or IN-vaccinated mice after lethal challenge (10xLD_50_) with six different influenza A virus challenge strains: **(C)** 1918 pandemic H1N1, **(D)** H7N9, **(E)** highly pathogenic avian H5N8, **(F)** chimeric avian H7N1, **(G)** chimeric avian H6N1, and **(H)** chimeric H10N7 virus. Error bars represent standard deviation. n.s; not significant

To evaluate vaccine efficacy against viral challenge, cohorts of BPL-inactivated vaccinated animals were challenged with six IAV strains at a 10x mouse LD_50_ dose (Supplementary Table 2). Mock-vaccinated animals steadily lost weight and showed 100% mortality following challenge with each of these viruses (Fig. 1C-H). IM and IN vaccination provided 100% protection against lethal 10x LD_50_ challenge with the fully reconstructed 1918 pandemic H1N1 and zoonotic H7N9 viruses (Fig. 1C,D) with little associated weight loss. Lethal 10x LD_50_ challenge with a highly pathogenic avian influenza (HPAI) H5N8 virus (Fig. 1E) showed 90% protective efficacy for this antigenically variant, systemically replicating HPAI virus following IM vaccination. Protective efficacy of IN vaccination against this HPAI virus was less than that observed for IM vaccination, with 70% survival following lethal H5N8 challenge.

To study the effects of vaccination following lethal challenge with homosubtypic, partially heterosubtypic (challenge with a virus with a novel HA subtype) and completely heterosubtypic (challenge with a virus with a novel HA and NA subtype), immunized mice were challenged with chimeric avian H7N1, H6N1, and H10N7 (*20, 22*), respectively. These viruses were chosen to represent different antigenic distance from the vaccine antigens (Supplementary Table 2). The H7N1 virus HA matched the vaccine H7 HA, along with a minor mismatch in N1. In contrast, the H6N1 challenge used the same NA as in the H7N1 challenge but in this case with an HA subtype not contained in the vaccine. The H10N7 virus expressed both HA and NA subtypes not contained in the vaccine. In all three challenge experiments, both IM- and IN-vaccinated mice showed 100% survival.

Vaccinated and PBS-vaccinated (mock) mice were challenged with a 10x LD_50_ dose of the H7N1 subtype virus. Vaccinated mice lost very little weight and all survived lethal challenge, in contrast to mock-vaccinated mice, which showed a rapid decline in body weight with 100% fatality occurring between days 6-8 post-challenge (Fig. 1F). A second cohort of vaccinated and mock-vaccinated mice were challenged with a 10x LD_50_ dose of the H6N1 virus. Vaccinated mice lost less weight and had 100% survival following lethal challenge, in contrast to mock-vaccinated, challenged mice, which showed a rapid decline in body weight and 100% fatality by days 6-7 post-challenge (Fig. 1G). Interestingly, IM vaccinated mice showed more early weight loss than IN vaccinated mice, with weight loss on days 1-4 weight loss similar to mock vaccinated groups, but with recovery and 100% survival. In a third cohort, mice were challenged with a 10x LD_50_ dose of the H10N7 virus to examine protection of vaccination against completely heterosubtypic viral infection. Both IM and IN vaccinated mice lost significantly less weight through day 4, but then rapidly recovered and had 100% survival following lethal challenge, in contrast to mock vaccinated challenged mice, which showed a steady decline in body weight and 100% fatality by days 6-7 post-challenge (Fig. 1H). Together, these results showed that vaccination of mice resulted in broad protection from a variety of lethal challenges with viruses of varying degrees of HA and NA antigenic distances. In five of the six different lethal viral challenge experiments, both IM and IN mice showed 100% survival, with 0% survival of mock-vaccinated animals. For HPAI H5N8 challenge, IM and IN vaccination afforded 90% and 70% protection, respectively.

### Pulmonary gene expression responses during mismatched and heterosubtypic challenge in mice

To determine the effects of vaccination on pulmonary inflammatory and immune responses during viral infection, expression microarray analysis was performed on lung tissue collected from mock and vaccinated animals on day 6 following challenge with chimeric H7N1, H6N1, and H10N7 chimeric viruses. Viral replication in whole lung tissue measured by qPCR did not detect M gene viral RNA by 3 to 6 days post-challenge with H7N1 and H6N1 viruses, and showed nearly 90% reduction in viral RNA by 6 days post-H10N7 challenge (Fig. 2A). Analysis of variance (ANOVA) identified significantly differentially expressed genes (≥2-fold difference in median expression, p<0.05) between mock, IN and IM challenged animals. Expression levels of genes associated with type I interferon (IFN) responses, lymphocyte activation, reactive oxygen species responses and DNA damage, and programmed cell death were significantly reduced in IN and IM vaccinated mice challenged with H7N1, H6N1, and H10N7 compared to PBS-vaccinated, challenged animals (Fig. 2A). Higher expression levels of these genes in PBS-vaccinated animals correlated with higher viral replication levels. Concordant with the weight loss in vaccinated animals observed following completely heterosubtypic H10N7 lethal challenge compared to either H7N1 and H6N1, IN- and IM-vaccinated animals showed slightly stronger immune-response related gene expression in response to H10H7 (Fig. 2A). These animals also showed slightly higher levels of viral replication.

**Figure 2.**
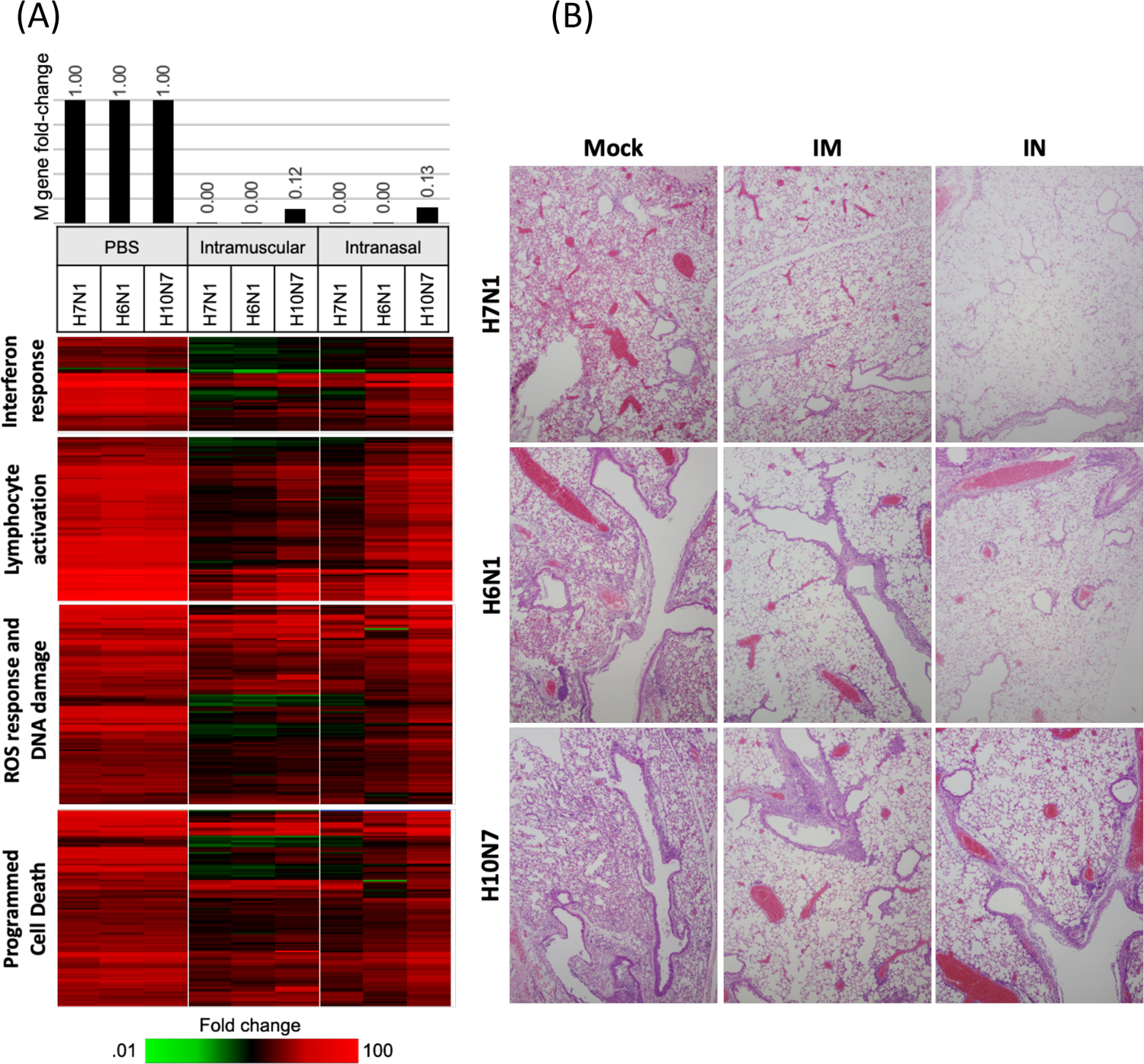
Quantitative RT-PCR for influenza viral RNA, lung inflammatory responses, and histopathology of vaccinated mice. **(A)** Top panel, bar graph showing relative expression of IAV M gene mRNA in vaccinated mouse lung compared to mock-vaccinated animals as measured by qRT-PCR. Lower panels, differences in lung gene expression responses in lungs of mock, IM, and IN vaccinated mice identified by ANOVA (≥2-fold difference in median expression level, p<0.01) on day 6 post-challenge with heat maps show the relative expression of type I interferon response, lymphocyte activation, ROS response and DNA damage, and programmed cell death. Genes with increased expression are show in red, genes with no change as black, and genes showing decreased expression in green. **(B)** Lung histopathology of mock-, IM-, and IN-vaccinated mice lethally challenged with chimeric avian H6N1, H7N1, or H7N10 viruses (10xLD_50_ dose), and analyzed at day 5 post-infection. In each case, mock-vaccinated animals showed a widespread, severe viral pneumonia with necrotizing bronchitis and bronchiolitis, alveolitis. In contrast, IM- or IN-vaccinated animals showed an absence of pneumonia, with no bronchitis, bronchiolitis, or alveolitis. Aggregates of lymphoid tissue were observed in peri-bronchiolar and peri-bronchiolar spaces in vaccinated animals. Original magnifications 20x. See Supplementary Figs 5-7 for additional pathological analyses.

IN vaccinated animals showed more robust gene expression responses compared to IM vaccinated mice; direct comparison (t test, 2-fold difference in median expression, p value <0.05) of IN- and IM-vaccinated animals revealed subtle but significant differences in gene expression (Supplementary Fig. S3A) that showed enrichment of pathways for neutrophil adhesion, IFN signaling, cytokine/chemokine signaling, dendritic cell maturation and other pathways involved in innate response to infection (Supplementary Fig. S3B).

### Lung pathology and immune cell infiltrates during mismatched and heterosubtypic challenge in mice

Histopathological examination was performed on mouse lung sections at day 5 post-viral challenge (Fig. 2B and Supplementary Fig. S4-6) for H7N1, H6N1, and H10N7 experiments.

Lung sections of mock-vaccinated mice challenged with H7N1 showed marked pathological changes involving over 50% of the lung parenchyma, including multifocal, moderate-to-severe, necrotizing bronchitis and bronchiolitis, along with moderate-to-severe alveolitis with pulmonary edema and fibrinous exudates (Fig. 2B and Supplementary Fig 5). Influenza viral antigen staining showed widespread positivity in respiratory epithelial cells and in alveolar epithelial cells. In contrast, lung sections from IM- and IN-vaccinated animals challenged with H7N1 (both HA and NA represented in the vaccine) showed minimal histopathological changes, an absence of alveolitis and no viral antigen in alveolar epithelial cells. The respiratory epithelium of bronchi and bronchioles was intact. Sections from mock-vaccinated mouse lungs showed occasional CD19+ B cells and CD3+ T lymphocytes, but abundant Ly6G+ neutrophils throughout the lung parenchyma. In contrast, IM- and IN-vaccinated mouse lungs showed increased CD19+ and CD3+ lymphocyte aggregates especially prominent in perivascular and peribronchiolar locations and a marked reduction in lung parenchymal neutrophils. Large aggregates of CD19-plasma cells were observed focally in the lungs of vaccinated mice.

Similarly, lung sections of mock-vaccinated mice challenged with H6N1 showed marked pathological changes involving over 50% of the lung parenchyma, including multifocal, moderate-to-severe, necrotizing bronchitis and bronchiolitis, along with moderate-to-severe alveolitis with pulmonary edema and fibrinous exudates (Fig. 2B and Supplementary Fig S5). Influenza viral antigen staining showed widespread positivity in respiratory epithelial cells and in alveolar epithelial cells. In contrast, lung sections, from IM- and IN-vaccinated animals challenged with the partially heterosubtypic H6N1 virus, showed minimal histopathological changes, and rapidly reproliferating respiratory epithelium in bronchi and bronchioles characterized by abundant mitotic figures, an absence of alveolitis and no viral antigen in alveolar epithelial cells. Sections from mock-vaccinated mouse lungs showed occasional CD19+ B cells, CD3+ T cells, and abundant Ly6G+ neutrophils throughout the lung parenchyma, with prominent parenchymal neutrophil infiltrates and neutrophil margination from pulmonary blood vessels. In contrast, IM- and IN-vaccinated mouse lungs showed markedly increased CD19+ B and CD3+ T lymphocyte aggregates, especially in perivascular and peribronchiolar locations and a marked reduction in lung parenchymal neutrophils. Lung sections from vaccinated mice showed foci of bronchiolar intra-epithelial CD19+ B cells, and multifocal accumulations of CD-19-plasma cells in peribronchiolar and perivascular locations. These results demonstrated that both IM and IN vaccination in mice resulted in complete protection from lethal challenge with a partially heterosubtypic lethal H6N1 viral challenge, associated with dramatic reductions in viral titer, pathologic changes, and host immune and inflammatory responses in lung, and a marked increase in B and T cell aggregates in the lungs of vaccinated mice.

Lung sections of mock-vaccinated mice challenged with H10N7 showed marked pathological changes involving over 50% of the lung parenchyma, including multifocal, moderate-to-severe, necrotizing bronchitis and bronchiolitis, along with moderate-to-severe alveolitis with a neutrophil-predominant, mixed cellularity inflammatory infiltrate, pulmonary edema and fibrinous exudates (Fig. 2B and Supplementary Fig S6). Influenza viral antigen staining showed widespread positivity in respiratory epithelial cells and in alveolar epithelial cells. In contrast, lung sections, from IM- and IN-vaccinated animals challenged with the completely heterosubtypic H10N7 virus, showed a marked reduction in histopathological changes, rapidly reproliferating respiratory epithelium in bronchi and bronchioles characterized by abundant mitotic figures, little-to-no alveolitis with little viral antigen in alveolar epithelial cells but viral antigen detected in alveolar macrophages multifocally. Sections from mock-vaccinated mouse lungs showed occasional CD19+ B and CD3+ T lymphocytes, and large infiltrates of Ly6G+ neutrophils throughout the lung parenchyma with prominent neutrophil margination from pulmonary blood vessels. In contrast, IM- and IN-vaccinated mouse lungs showed markedly increased CD19+ and CD3+ lymphocyte aggregates, especially in perivascular and peribronchiolar locations and a marked reduction in lung parenchymal neutrophils. These results demonstrated that both IM and IN vaccination in mice resulted in complete protection from lethal challenge with a completely heterosubtypic H10N7 virus that was associated with dramatic reductions in viral titer, pathologic changes, and host immune and inflammatory responses in lung.

Immunization with lower doses (1/4 antigen) of the same vaccine provided 100% protection against 10x LD_50_ lethal challenge with completely heterosubtypic H10N7 and partially heterosubtypic H6N1 viruses in mice (Supplementary Fig S7), suggesting that a lower dose of the vaccine could still provide a high level of protection. Passive serum transfer experiments in mice, in which serum from vaccinated animals was injected intraperitoneally 1 day prior to challenge in unvaccinated mice with intrasubtypic H7N1, provided 100% protection with serum from IM-vaccinated, but not from IN-vaccinated animals (Supplementary Fig. S8A), consistent with the lower levels of serum anti-viral antibody observed in IN-vaccinated animals (Fig. 1A-B). Serum transfer experiments followed by H6N1 challenge, in which the HA subtype was not contained in the vaccine, while the N1 NA subtype was, produced analogous results to intrasubtypic H7N1 challenge, in that serum from IM-vaccinated mice saved unvaccinated H6N1-challenged while serum from IN-vaccinated mice did not (Supplementary Fig. S8B). In this case, protection was likely afforded by anti-neuraminidase antibody in vaccinated serum, and/or anti-group 1 HA stalk antibody. In contrast, passive serum transfer from either IM- or IN-immunized mice provided no protection against heterosubtypic H10N7 challenge in unvaccinated mice (Supplementary Fig. S8C), suggesting strongly that the complete heterosubtypic protection observed (Fig. 1H and Supplementary Fig. S6) is not primarily mediated by serum antibodies, but is likely mediated by cellular immune responses. Having demonstrated broad protective immunity in vaccinated mice following a variety of lethal challenge experiments including challenge with completely heterosubtypic viruses, the effects of vaccination against mismatched and heterosubtypic IAV viral challenge was next evaluated in ferrets.

### Immune responses to IM and IN vaccination in ferrets and protection against mismatched and heterosubtypic viral challenge

Ferrets were primed and boosted 28 days later by IM and IN vaccination with 400ug total protein (100 ug per subtype). IN vaccination was performed without adjuvant, and IM immunization was performed using a squalene-based adjuvant (Supplementary Fig. S1B). Mock-immunized control animals were intranasally inoculated with PBS or intramuscularly inoculated with PBS including adjuvant. The sequence similarities between the vaccine strains and the challenge strains are summarized in Supplementary Table 2. As shown in Fig. 3A,B, the vaccine was highly immunogenic in ferrets and both IN and IM immunization elicited significant serum IgG antibody responses to homologous HAs (H1, H3, H5, H7) and NAs (N1, N3, N8, N9) as well as serum group-1 and group-2 HA stalk antibodies and hemagglutination inhibition (HAI) antibodies (Supplementary Fig. S9A,B). Similar to what was observed in mice, IM immunization induced generally higher serum IgG antibody responses than IN immunization in ferrets (Fig. 3A,B).

**Figure 3.**
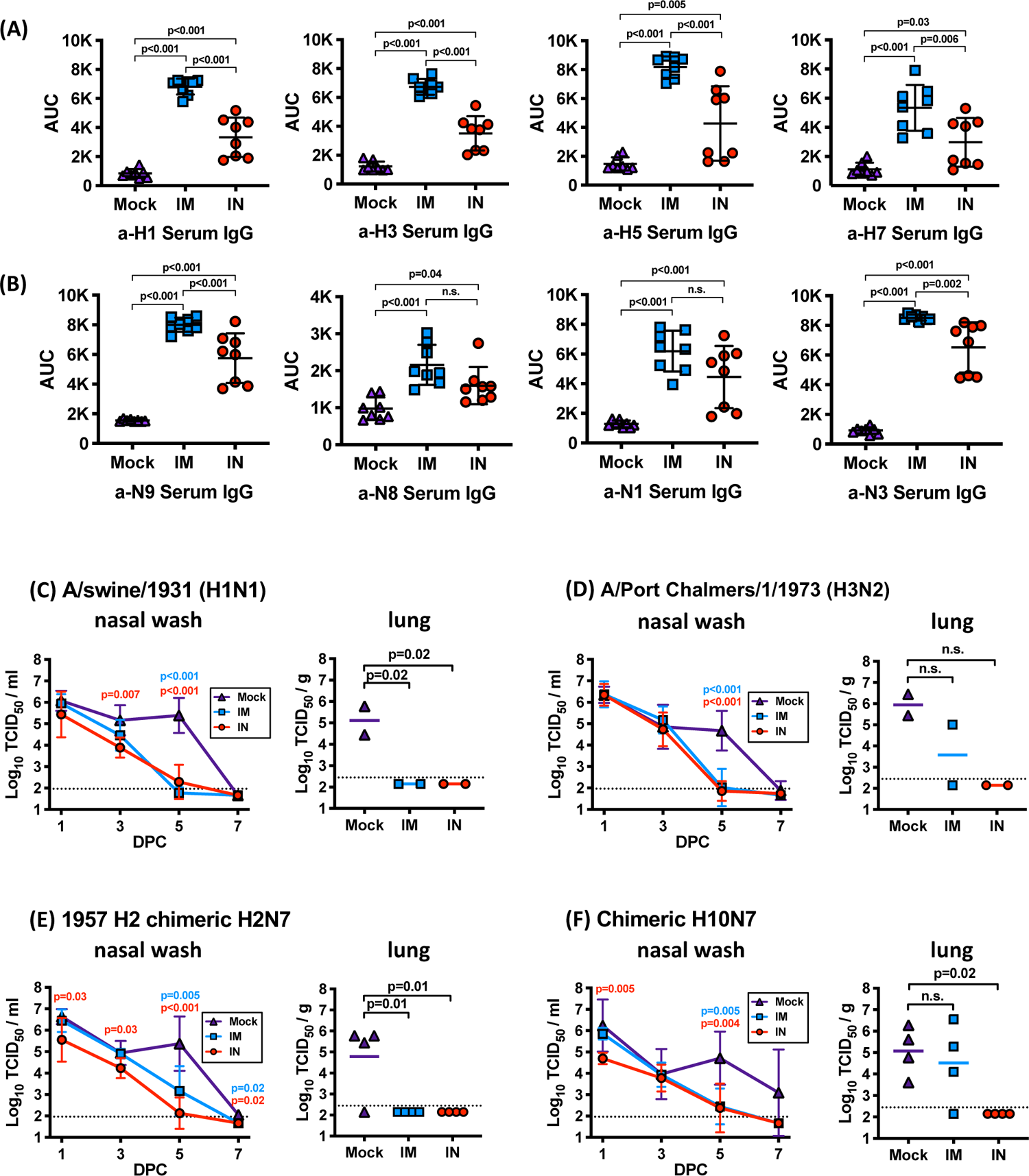
Immunogenicity and reduction in viral titers of BPL-inactivated vaccinated ferrets. Serum IgG levels against (**A**) four vaccine hemagglutinin (HA) antigens and (**B**) four vaccine neuraminidase (NA) antigens were measured 3 weeks after the boost immunization in mock-, IM-, or IN-vaccinated mice using ELISA. Ordinary ANOVA test and post hoc Tukey’s multiple comparison test were used to compared antibody levels between groups. Error bars represent standard deviation. **(C-F).** Reductions in nasal wash and lung viral titers in IM- or IN-vaccinated ferrets as compared to mock-vaccinated ferrets. Nasal wash titers were measured at days 1, 3, 5, and 7 following challenge, and lung titers were determined on day 5 following challenge. (C). Reductions in titers following A/swine/1931 (H1N1) challenge. (D). Reductions in titers following A/Port Chalmers/1/1973 (H3N2) challenge. (E). Reductions in titers following chimeric 1957 pandemic H2N7 challenge. (F). Reductions in titers following chimeric avian H10N7 challenge. Ordinary ANOVA test and post hoc Dunnett’s multiple comparison test were used to compare viral titers in IM- or IN-vaccinated ferrets to mock-vaccinated ferrets. Error bars represent geometric standard deviation. n.s; not significant

The effect of vaccination on homosubtypic A/H1N1 viral challenge in ferrets was next evaluated. Adult female ferrets were primed and boosted as above with the tetravalent, whole virus BPL-inactivated vaccine. Vaccine efficacy was evaluated following mismatched, intrasubtypic challenge with A/swine/1931 H1N1 virus (Supplementary Table 2). HAI assay demonstrated a statistically significant, approximately 4-fold difference in HAI titers between the vaccine H1 HA and the challenge H1 viruses (Supplementary Fig. S10A). The N1 NA shared only 85.5% nucleotide identity with the N1 NA in the vaccine, demonstrating that the challenge virus has substantial antigenic difference compared to vaccine antigens. Viral titers in ferret nasal wash in IM- and IN-vaccinated ferrets were significantly reduced to near undetectable levels by day 5 post-challenge as compared to mock-vaccinated ferrets (Fig. 3C), and lung titers collected from IM- and IN-vaccinated ferrets at day 4 post-challenge showed no detectable levels of viral replication.

Similarly, vaccinated and mock-vaccinated ferrets were challenged with a partially heterosubtypic human seasonal IAV, A/Port Chalmers/1973 (H3N2) (Supplementary Table 2). In this case, the H3 was antigenically mismatched to the avian H3 HA in the vaccine, as supported by cross-HAI evaluation (Supplementary Fig. S10B) and sequence identity to the vaccine H3 HA (83.8%), while the challenge virus expressed an N2 subtype NA not contained in the vaccine. The closest vaccine NA sequence shared only 43.5% identity with the challenge virus N2 subtype. Post-challenge viral titers in nasal wash in IM- and IN-vaccinated ferrets were significantly reduced to near undetectable levels by day 5 post-challenge (Fig. 3D), and lung titers collected at day 5 post-challenge showed no detectable levels of viral replication in the IN-vaccinated group.

To measure vaccine protective efficacy in ferrets against completely heterosubtypic IAV challenge, two cohorts of vaccinated and mock-vaccinated ferrets were challenged with either A/H2N7 or A/H10N7 (Supplementary Table 2). Sequence identity of the challenge HA and NA subtypes as compared to the vaccine components ranged from 44.9-53.9%. Viral titers in nasal wash in IM- and IN-vaccinated A/H2N7-challenged ferrets were significantly reduced by day 5 post-challenge (Fig. 3E), and lung titers collected at day 5 post-challenge showed no detectable levels of viral replication in both the IM- and IN-vaccinated groups. Similarly, challenge viral titers in ferret nasal wash in IM- and IN-vaccinated A/H10N7-challenged ferrets were significantly reduced by day 5 post-challenge (Fig. 3F). Lung titers collected at day 5 post-challenge showed no detectable levels of viral replication in IN-vaccinated animals, while IM-vaccinated animals had viral titers that were not significantly reduced compared to mock-vaccinated animals.

### Host gene expression responses and qRT-PCR for viral RNA during mismatched and heterosubtypic challenge in ferrets

To characterize the host gene expression response to viral challenge in PBS- and vaccinated mice, RNA expression microarray analysis and IAV M gene qRT-PCR was performed on lung tissue collected on day 5 post-challenge. ANOVA was performed to identify significantly differentially expressed genes (≥2-fold difference in median expression, p<0.05) between mock, IN and IM challenged animals. Due to the small number of animals, the analysis was performed with all 4 viruses in each group (mock-, IM- and IN-vaccinated). These sequences were enriched for pathways associated with the innate antiviral response, including IFN signaling, cytokine signaling, lymphocyte activation, and oxidative damage DNA repair responses (Fig, 4A) which were highly expressed in control animals but significantly lower in the IM- and IN-vaccinated animals (Fig. 4A), consistent with little viral replication in the lungs of these animals. No significant differences in lung gene expression responses were identified between IM- and IN-vaccinated, challenged ferrets.

**Figure 4.**
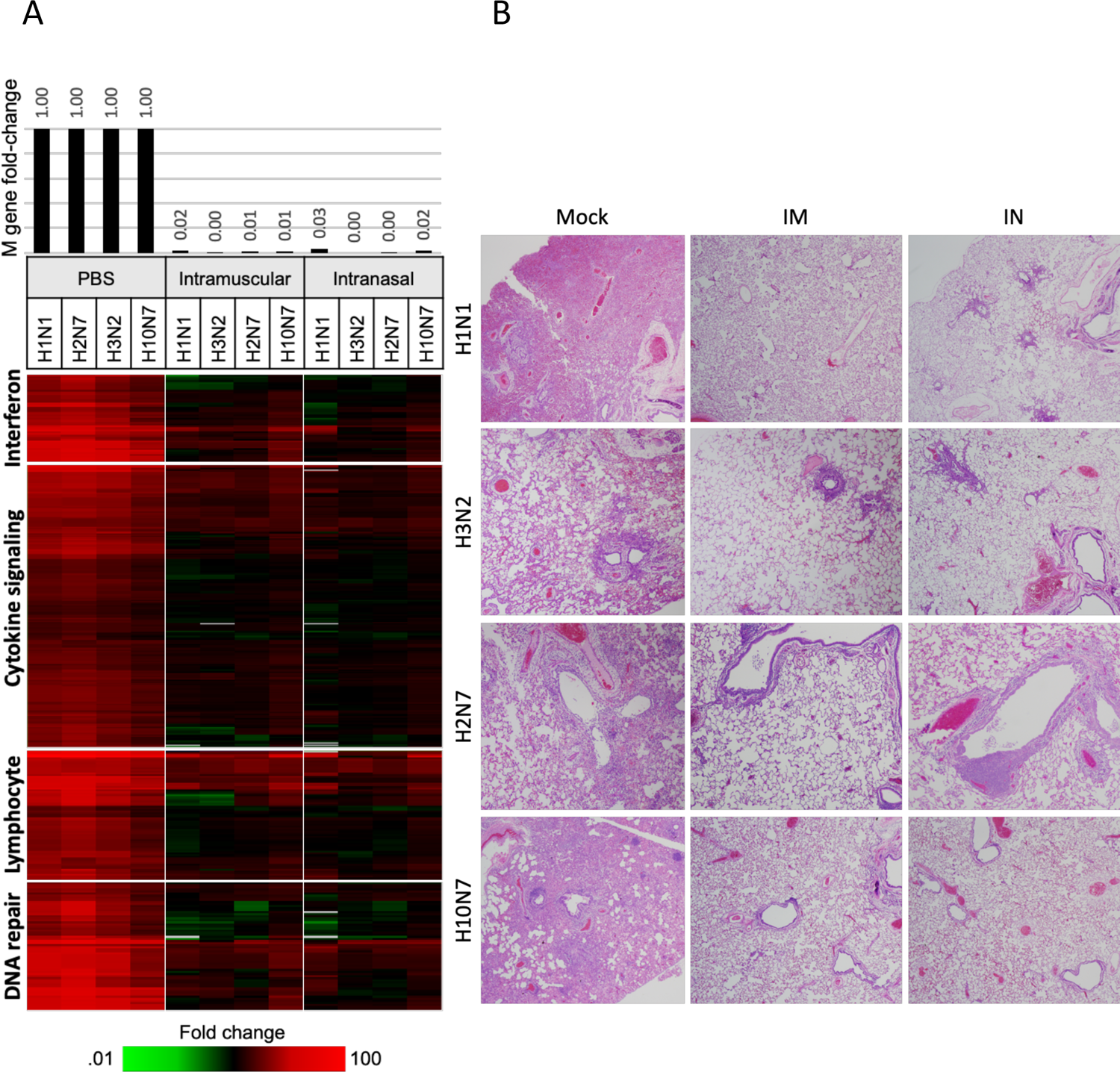
Quantitative RT-PCR for influenza viral RNA, lung inflammatory responses, and histopathology of vaccinated ferrets. **(A)** Top panel, bar graph showing relative expression of IAV M gene mRNA in vaccinated ferret lung compared to mock-vaccinated animals as measured by qRT-PCR. Lower panels, differences in lung gene expression responses in lungs of mock, IM, and IN vaccinated ferrets identified by ANOVA (≥2-fold difference in median expression level, p<0.01) on day 5 post-challenge with heat maps show the relative expression of type I interferon responses, cytokine signaling, lymphocyte activation, and DNA repair. **(B)** Lung histopathology of mock-, IM-, and IN-vaccinated mice lethally challenged with A/swine/1931 (H1N1), A/Port Chalmers/1/1973 (H3N2), or completely heterosubtypic challenge with chimeric H2N7 or H10N7 challenge viruses. In each case, mock-vaccinated animals showed a widespread, severe viral pneumonia with necrotizing bronchitis and bronchiolitis, alveolitis. In contrast, IM- or IN-vaccinated animals showed an absence of pneumonia, with no bronchitis, bronchiolitis, or alveolitis. Aggregates of lymphoid tissue were observed in peri-bronchiolar and peri-bronchiolar spaces in vaccinated animals. Original magnifications 20x. See Supplementary Figs S11-14 for additional pathological analyses.

### Lung pathology during mismatched and heterosubtypic viral challenge in ferrets

Histopathological analysis was performed on ferret lung sections at day 5 post-viral challenge (Fig. 4B and Supplementary Fig. S11-14). Lung sections of mock-vaccinated ferrets challenged with swine/Iowa/1931 (H1N1) virus showed marked pathological changes involving over 50% of the lung parenchyma, including multifocal, moderate-to-severe, necrotizing bronchitis and bronchiolitis (Fig. 4B and Supplementary Fig. S11), along with moderate-to-severe alveolitis with a neutrophil-predominant, mixed inflammatory cell infiltrates, pulmonary edema and fibrinous exudates, a pathology remarkably similar to that seen with ferret infection with the 1918 pandemic H1N1 virus (*23*). Influenza viral antigen staining showed widespread positivity in respiratory epithelial cells and in alveolar epithelial cells. In contrast, lung sections from IM- and IN-vaccinated animals showed minimal histopathological changes, including mild, focal bronchiolitis, an absence of alveolitis and no viral antigen in alveolar epithelial cells.

Histopathological analysis was performed on ferret lung sections at day 5 post-viral challenge with the partially heterosubtypic A/Port Chalmers/1973 (A/H3N2) virus (Fig. 4B and Supplementary Fig S12). Lung sections of mock-vaccinated ferrets showed marked pathological changes involving over 50% of the lung parenchyma, including multifocal, moderate-to-severe, necrotizing bronchitis and bronchiolitis, along with moderate-to-severe alveolitis with a mixed inflammatory cell infiltrate, and focal pulmonary edema and fibrinous exudates. Influenza viral antigen staining showed widespread positivity in respiratory epithelial cells and in alveolar epithelial cells. In contrast, lung sections from IM- and IN-vaccinated animals showed no pneumonia, an absence of alveolitis, and no viral antigen in alveolar epithelial cells, while also showing multi-focal peri-bronchiolar lymphoid infiltrates.

Lung pathology from fully heterosubtypic viral challenges (with A/H2N7 and A/H10N7) were evaluated next. Lung sections of mock-vaccinated H2N7 challenged ferrets showed marked pathological changes (Fig. 4B and Supplementary Fig. S13) involving over 50% of the lung parenchyma, including multifocal, moderate-to-severe, necrotizing bronchitis and bronchiolitis, along with moderate-to-severe alveolitis with a mixed inflammatory cell infiltrate and numerous intra-alveolar inflammatory cells. Influenza viral antigen staining showed widespread positivity in respiratory epithelial cells and in alveolar epithelial cells. Lung sections of mock-vaccinated H10N7 challenged ferrets showed marked pathological changes Fig. 4B and Supplementary Fig. S14), involving over 50% of the lung parenchyma, including multifocal, moderate-to-severe, necrotizing bronchitis and bronchiolitis, along with moderate-to-severe alveolitis with a neutrophil-predominant, mixed inflammatory cell infiltrate, numerous intra-alveolar inflammatory cells, and widespread pulmonary edema and fibrinous exudates. Influenza viral antigen staining showed widespread positivity in respiratory epithelial cells and in alveolar epithelial cells. In contrast, lung sections from IM- and IN-vaccinated animals challenged with H2N7 showed no pneumonia, bronchitis or bronchiolitis, but also showed prominent peri-bronchiolar lymphoid nodules. No alveolitis was noted and no influenza viral antigen was detected in alveolar epithelial cells in vaccinated ferrets (Supplementary Fig. S14). Similarly, lung sections from IM- and IN-vaccinated animals challenged with H10N7 showed no pneumonia, no bronchitis, bronchiolitis, or alveolitis. Peri-bronchiolar lymphoid nodules were observed. No influenza viral antigen was detected in alveolar epithelial cells, but as with H3N2 challenge above, some viral antigen was detected in bronchiolar respiratory epithelial cells in the absence of inflammation or histopathologic changes in IM-challenged animals, which may correspond to the detectable viral titers in 3 of 4 those animals (Fig. 3F). Similar to mouse studies, immunization with lower doses (1/4 antigen) provided protection against partially-heterosubtypic H3N2 and mismatched H1N1 challenge in ferrets (Supplementary Fig S15), suggesting that a lower dose of the multivalent vaccine candidate could still provide a high level of protection.

### GMP manufacture, toxicology and immunogenicity studies

The systemic toxicity, local tolerance, and immunogenicity of the BPL influenza vaccine (administered IN or IM) was evaluated in New Zealand white rabbits. No mortality was observed following administration of the vaccine IN or IM. There were no clinical observations, injection or instillation site observations, changes in body weights, changes in food consumption, changes in body temperatures, systemic toxicity, local tolerance or ocular effects attributed to administration of the vaccine by either route. Rabbits that received the vaccine intranasally or intramuscular mounted serum antibody responses against the vaccine HA and NA antigens on day 45 (Supplementary Fig. S16).

## Discussion

A challenge to development of a “universal” influenza vaccine is the elicitation of effective broadly neutralizing antibodies and memory T cell responses. Currently, annual influenza virus vaccines are produced each year based on surveillance of circulating strains and predictions of which strains will be circulating the following season (*24*). While this approach can yield limited success, in many years strain-match predictions are imperfect and seasonal vaccines can be of limited effectiveness. Development of a “universal” influenza vaccine could be used as a super-seasonal vaccine that provides protection against new seasonal strains without the need for predictive antigenic matching, as well as provide protection against newly emerging pandemic and zoonotic influenza virus infections (*7*). Efforts to create a universal influenza vaccine have been ongoing for over four decades (*12*), while in the last five years this idea has gained renewed impetus from funders (*7*). Numerous strategies to achieve this goal are being currently pursued, including vaccines based on hemagglutinin head or stalk antigens, neuraminidase, the M2 protein exodomain, live attenuated influenza vaccines, and T cell-based vaccines targeting viral peptide epitopes, as recently reviewed (*25, 26*). Several vaccine candidates have advanced into early clinical development (*27–29*), but whether these strategies will induce broad protection in humans has not yet been determined.

The approach taken in this study was to develop a broadly protective vaccine using beta-propiolactone (BPL) inactivated whole avian influenza viruses that contain full-length, properly folded HA and NA proteins, as well as other viral proteins (such as NP and M proteins) that have been shown to have conserved T cell epitopes (*30*). BPL-inactivated vaccines also have the advantage that they are simple and cheap to manufacture and have reduced cold-chain requirements. This study demonstrates that a broadly protective influenza vaccine, comprised of four BPL-inactivated whole low pathogenicity avian viruses, demonstrated near-universal protection from lethal viral infection against homologous, partially heterosubtypic, and completely heterosubtypic viral challenge in both mice and ferrets. For this vaccine, the four IAVs were chosen because they represent a broad consensus of the cladal HA and NA distribution of influenza A viruses. Avian IAV HA proteins also have low levels of glycosylation on the HA head as compared to human viruses, which allow for development of antibody responses against protein antigens that might be masked in seasonal IAVs. Moreover, because whole virions are used that likely provide T cell epitopes, vaccination should promote the development of broader memory T cell responses compared to purified vaccine antigens. The inactivated vaccine, delivered either IM or IN, is likely to be safe in humans as no toxicity was observed in mice or ferrets or in a rabbit toxicity study conducted as part of GMP manufacture.

In a prior study utilizing a vaccine consisting of a cocktail of four viral-like particles expressing the four HA proteins used here (*20*), broad protection against mismatched and completely heterosubtypic challenge was observed in mice. In this follow-up study, broad and potent protective efficacy was observed for a BPL-inactivated whole virus vaccine both in mouse and ferret model. Advantages of the current vaccine include ease of production, immunization with four HA antigens which induce systemic and respiratory antibodies against the HA heads and stalks, the addition of four divergent NA proteins which also induce systemic and respiratory antibodies, and internal viral proteins which likely serve as targets of T cell responses. Supporting this conclusion, the vaccine provided near 100% protective efficacy against mismatched or completely heterosubtypic challenge in mice and ferrets. In mice, vaccinated animals were protected against mismatched lethal challenge with pathogenic 1918 pandemic H1N1, H7N9, HPAI H5N8, and chimeric avian H7N1 virus challenges. Significantly, complete protection was afforded following challenge with the H6N1 virus, expressing a heterosubtypic HA (H6) not contained in the vaccine, and even more significantly, complete protection was observed following challenge with the fully heterosubtypic H10N7 virus, expressing both HA and NA subtypes not contained in the vaccine. Similarly, significant protection was observed in ferrets following completely heterosubtypic challenge with H2N7 or H10N7 viruses. These results are consistent with the desired characteristics of a “universal” influenza vaccine that could be of value in human vaccination programs.

Protective efficacy of vaccine-induced antibodies was evaluated in passive transfer of serum from mock-, IM-, or IN-vaccinated mice to naïve (unvaccinated) mice 1 day prior to lethal challenge with mismatched H7N1, partially heterosubtypic H6N1, or completely heterosubtypic H10N7 (Supplementary Fig. 9). Serum from IM-vaccinated mice afforded complete protection from H7N1 and partially heterosubtypic H6N1 virus lethal challenge, but not from completely heterosubtypic H10N7 challenge. Serum antibodies mediating survival following H6N1 challenge might include HA stalk antibodies, but protection was more likely provided by cross-reactive NA antibodies from the vaccine N1 antigen, since no serum protection was seen following completely heterosubtypic H10N7 challenge. These results demonstrate that a protective antibody response was generated against homologous IAV but suggest that other vaccine-induced immune mechanisms involving cellular immunity must be involved in protection against lethal challenge with heterosubtypic viruses.

In both IM- and IN-vaccinated mice and ferrets peri-bronchiolar and peri-vascular lymphoid aggregates were observed, and in mice it was possible to perform immunostaining to show that these aggregates contained both CD19+ B and importantly, CD3+ T lymphocytes. That IN-vaccinated mice and ferrets demonstrated potent protective efficacy even against completely heterosubtypic challenge despite mounting lower levels of serum antibody responses suggests that mucosal immunity plays an important role in anti-influenza virus responses, both through humoral and cellular immunity. As a further metric of vaccine effectiveness, vaccination was associated with a significant reduction in lung pathology and inflammatory gene expression responses in both mice and ferrets. The reduction in lung neutrophils during lethal challenge resulting from vaccination is likely critical to protection against lethal infection, as neutrophils and the ROS molecules they produce and excrete are major contributors to severe pulmonary pathology during fatal IAV infections (*22, 31–33*). Also concordant with reductions in viral replication and lung neutrophils, expression microarray analysis showed significant reductions in expression of genes in type I IFN, ROS damage, and cell death pathways compared to mock-vaccinated and challenged mice and ferrets. In mice, increasing activation of these genes and associated pathways was observed with partially and completely heterosubtypic viral challenge in mice.

Further, the route of vaccination also affected lung host gene expression responses in mice. While both IM and IN vaccination were protective against lethal challenge in mice, expression analysis showed lower expression of inflammatory gene expression response pathways correlated with IM vaccination compared to IN vaccination. This vaccination route-dependent effect suggests that in mice mucosal or systemic vaccination affects molecular aspects of host immune responses, which arise from differences in effector functions or epithelial transport of IgA and IgG antibodies (*34*), or even a larger pool of adaptive immune cells in the periphery at vaccination. While both IM and IN vaccination were protective against lethal challenge in mice and ferrets, further studies are underway to better understand variables associated with the route of vaccination.

In summary, these preclinical studies in mice and ferrets demonstrate that a BPL-inactivated vaccine possesses many features of a successful universal influenza vaccine, with the potential to offer protection in human populations as a supra-seasonal and pre-pandemic IAV vaccine. In additional studies, humoral and cellular immune correlates of protection will be evaluated, as will vaccine efficacy in animals with a pre-existing exposure history to influenza A viruses to better mimic the complex immune repertoire exposure history to influenza viruses in humans, and whether vaccination results in reduction in transmission of influenza in ferrets.

## Acknowledgements

This work was supported in part by the Intramural Research Programs of the National Institutes of Health and National Institute of Allergy and Infectious Diseases (NIAID) and in part by a grant from the Bill and Melinda Gates Foundation (OPP1178956). Animal care was performed by the Comparative Medicine Branch, NIH/NIAID. The rabbit toxicology study was performed under a contract from Battelle, with support of the Division of Microbiology and Infectious Diseases, NIAID.

## Author contributions

JP, LMS, KAW, MJM, JCK, and JKT conceived and designed the study. JKP, SF, LMS, ZMS, AF, LM, YX, MR, NB, LQ, LAR, SW, KS, MG, KAW, JKT generated the laboratory data. JKP, LMS, MR, NB, LQ, LAR, KS, MG, IB, DMM, KAW, MJM, JCK, and JKT interpreted the data. JKP, DMM, KAW, MJM, JCK, and JKT wrote the manuscript. All authors critically reviewed the paper and approved of the final version of the paper for submission.

## Data and materials availability

The data and materials that support the findings of this study are available from the corresponding author upon reasonable request.

## Supplementary Materials

### Materials and Methods

#### Quadrivalent vaccine design and construction

Low pathogenicity avian viruses A/mallard/Ohio/265/1987 (H1N9), A/pintail/Ohio/339/1987 (H3N8), A/mallard/Maryland/802/2007 (H5N1), and A/Environment/Maryland/261/2006 (H7N3) were grown in MDCK cells followed by inactivation using β-propiolactone (BPL; catalog no. P5648; MilliporeSigma, USA). Viral culture supernatant was buffered with HEPES (catalog no. 15630; ThermoFisher, USA) at a final concentration of 0.1M and BPL was added (0.1% final). After overnight incubation at 4°C for the viral inactivation, BPL was hydrolyzed at 37°C for 90 min. Inactivated viruses were concentrated by ultracentrifugation at 50,000xg for 2h and purified using a 20-60% (w/v) discontinuous sucrose density gradient purification (100,000xg, 2h). A band at the 20-60% sucrose interface containing purified viruses was collected and the sucrose was removed by pelleting viruses (50,000xg, 2h) followed by virus resuspension in PBS. The total protein amount of the purified viruses was quantified using a bicinchoninic acid (BCA) protein assay kit (catalog no. 23225; ThermoFisher). For intranasal immunization in mice, 1 dose of vaccine was formulated to contain 1.5ug of each antigen (6ug total) in 50ul PBS. For intramuscular immunization in mice, 1 dose of vaccine was formulated to contain 1.5ug of each antigen (6ug total) in 25ul PBS and supplemented with 25ul of adjuvant AddaVax (catalog no. vac-adx-10; InvivoGen, USA). For the ferret study, 100ug of each antigen (400ug total) in 1ml PBS was used for 1 dose of intranasal immunization. For intramuscular immunization in ferrets, 100ug of each antigen (400ug total) in 250ul PBS was supplemented with 250ul of AddaVax (catalog no. vac-adx-10; InvivoGen). Low dose vaccines are prepared as described above, but with 1/4 antigen (i.e. 1.5ug total for mice, 100ug total for ferrets).

#### Challenge viruses

Fully reconstructed 1918 pandemic H1N1 virus was generated using a 12-plasmid reverse genetics system as previously described (*1, 2*) and passaged in MDCK cells. Avian influenza viruses (H6N1, H7N1, and H10N7) and recombinant H2N7 virus containing the 1957 H2 pandemic HA were generated as previously described (*1, 3*). A/Swine/Iowa/1931 (H1N1), A/Port Chalmers/1/1973 (H3N2), A/chicken/Netherlands/EMC-3/2014 (H5N8), and A/Shanghai/1/2013 (H7N9) viruses were passaged in embryonated specific-pathogen-free (SPF) chicken eggs (catalog no. 10100329; Charles River, USA). Supplementary Table 2 summarizes the challenge viruses used in this study and the hemagglutinin (HA) and neuraminidase (NA) identity to vaccine virus components. All viruses and infectious samples were handled under enhanced biosafety level 3 (BSL-3) laboratory conditions except for A/Swine/Iowa/1931 (H1N1) A/Port Chalmers/1/1973 (H3N2) which were handled in BSL-2 laboratory conditions. Experiments with the fully reconstructed 1918 pandemic virus and the highly pathogenic avian influenza (HPAI) H5N8 virus were conducted in accordance with the select agent guidelines of the National Institutes of Health (NIH), the Centers for Disease Control and Prevention and the United States Department of Agriculture, under the supervision of the NIH Select Agent and Biosurety Programs and the NIH Department of Health and Safety.

#### Mouse studies

Seven-to-eight-week-old female BALB/C mice (Jackson Laboratories, USA) were lightly anesthetized with isoflurane supplemented with O_2_ (1.5 L/min) before immunization or virus challenge. Mice were intranasally or intramuscularly immunized twice 4 weeks apart with the quadrivalent vaccines prepared as described above. Mock-vaccinated control mice received PBS or adjuvant without antigen. Serum samples were collected 3 weeks after the boost immunization to measure antibody responses elicited by the immunization. Lethal challenge infections (10x LD50 dose in 50ul inoculum per animal) were performed 4 weeks after the boost immunization. Post-challenge body weight and survival were monitored for 14 days from 5 mice/experimental condition. Ten mice, instead of 5, were used for the HPAI H5N8 challenge study. Mice were humanely euthanized if more than 25% of initial body weight was lost. For measuring viral loads and transcriptomics from H7N1, H6N1, and H10N7 challenge groups, lungs were harvested at day 6 (n=4) post-infection and immediately frozen in dry ice and stored at −80°C until processed. For histopathology, lungs were harvested at day 5 post-infection (n=2) followed by inflation and fixation using 10% neutral buffered formalin (NBF).

#### Ferret studies

Five-to-seven-month-old female ferrets (Triple F Farms, USA) were lightly anesthetized with isoflurane supplemented with O_2_ (1.5 L/min) before immunization or virus challenge. Ferrets were intranasally or intramuscularly immunized twice 4 weeks apart with the quadrivalent vaccines prepared as described above. Mock-vaccinated control ferrets received PBS or the adjuvant. Serum samples were collected 3 weeks after the boost immunization to measure antibody responses elicited by the immunization. Challenge infections (1ml inoculum per animal) were performed 4 weeks after the boost immunization. For A/swine/1931 (H1N1) and A/Port Chalmers/1973 (H3N2) challenge, 1×10^7^ plaque-forming unit (PFU) of each virus was used. For chimeric H2N7 and H10N7 challenge, 2×10^5^ PFU of each virus was used. To measure viral shedding in the upper respiratory tract following infection, nasal wash samples were collected at days 1, 3, 5, and 7 post-infection using 1ml of PBS. To measure viral shedding, transcriptomics, and histopathology in the lower respiratory tract, ferrets were euthanized at day 5 post-challenge and lungs were harvested. Left cranial lobes of the harvested lungs were immediately frozen in dry ice and stored at −80°C until processed for measuring viral loads and transcriptomics. For histopathology, remaining lungs were inflated and fixed using 10% NBF. All experimental animal work was performed in accordance with United States Public Health Service (PHS) Policy on Humane Care and Use of Laboratory Animals in an ABSL2 laboratory or an enhanced animal BSL3 (ABSL-3+) laboratory (for the viral challenge) as necessary at the National Institute of Allergy and Infectious Diseases (NIAID) of the NIH following approval of animal safety protocols by the NIAID Animal Care and Use Committee.

#### RNA isolation and expression microarray analysis

Frozen lungs, collected as described above, were lightly defrosted, homogenized in Trizol (catalog no. 15596018; ThermoFisher), and total RNA was isolated following manufacturer’s protocol. Isolated total RNA was purified using RNeasy Mini Kit (catalog no. 74106; Qiagen, Germany). Gene expression profiling experiments were performed using Agilent Mouse Whole Genome 44K microarrays (catalog no. G4122F; Agilent, USA). Ferret expression microarray analysis was performed using custom microarrays from Agilent Technologies (*4*). Fluorescent probes were prepared using Agilent QuickAmp Labeling Kit (catalog no. 5190-2305; Agilent) according to the manufacturer’s instructions. Each RNA sample was labeled and hybridized to individual arrays. Spot quantitation was performed using Agilent’s Feature Extractor software and all data were uploaded into Genedata Analyst 9.0 (Genedata, Switzerland). Data normalization was performed in Genedata Analyst 9.0 (Genedata) using central tendency followed by relative normalization using pooled RNA from mock infected mouse lung (n=4) or ferret lung (n=3) as a reference. Transcripts showing differential expression (2-fold, p< 0.01) between infected and control animals were identified by standard t test. The Benjamini-Hochberg procedure was used to correct for false positive rate in multiple comparisons. Panther and Ingenuity Pathway Analysis (IPA) was used for gene ontology and pathway classification [A-B].

#### Viral load determination in mouse and ferret lungs

Influenza viral titers in mouse and ferret lungs were quantified using qPCR from the RNA samples prepared as described above. Reverse transcription of total RNA was performed using the Superscript III first-strand cDNA synthesis kit (catalog no. 18080051; ThermoFisher) primed with an equal mix of oligo(dT) and the Uni12 influenza A specific primer: 5’ AGCRAAAGCAGG 3’. The IAV matrix gene amplicon was quantified using the following primers and probe sequences: forward primer, 5’-ARATGAGTCTTCTRACCGAGGTCG-3’; reverse primer, 5’-TGCAAAGACATCYTCAAGYYTCTG-3’; probe, 5’-[6-FAM] TCAGGCCCCCTCAAAGCCGA [BHQ1]-3’ (*5, 6*). Real-time PCR was performed on a Bio-Rad CFX384 Touch Real-Time PCR Detection System with TaqMan 2X PCR Universal Master Mix using a 10uL total reaction volume in duplicate. Ct values were normalized to the calibrator gene mouse GAPDH (catalog no. 4352932E; ThermoFisher).

#### Viral load determination in ferret nasal wash

Viral loads of ferret nasal wash samples were measured using 50% Tissue Culture Infectious Dose (TCID_50_) assay in MDCK cells. Reed and Muench method (*7*) was used for the TCID_50_ calculation.

#### Immunogenicity

Serum samples collected approximately three weeks post-boost immunization were used to investigate the immunogenicity of the quadrivalent vaccine. Hemagglutination inhibition (HAI) assay was performed as previously described (*8*). Enzyme-linked immunosorbent assay (ELISA) was also used to measure antibodies recognizing homologous HAs (H1, H3, H5, H7) and NAs (N1, N3, N8, N9) using recombinant HA and NA proteins. Antibodies recognizing group1 and group2 HA stalk were also measured. Recombinant proteins for the ELISA were designed based on previously published HA (*9*), NA (*10*), group 1 HA stalk (*11*), and group2 HA stalk (*12*) constructs with a Strep-Tag II affinity tag. Particularly, for the group 2 HA stalk, a chimeric HA consisting of a globular head of H4 HA and a stalk of H3 HA (cH4/3) was used. The recombinant proteins were expressed in insect cells, purified using Strep-Tactin Sepharose (catalog no. 2-1201; IBA GmbH, Germany), and quantified using BCA protein assay kit (catalog no. 23225; ThermoFisher) as previously described (*11*). Purified proteins were diluted in PBS (1µg/ml) and added to 96-well ELISA plates (50µl/well) (catalog no. 456537; ThermoFisher). The plates were incubated overnight at 4°C followed by the addition of blocking buffer (1% BSA in PBS, 100µl/well). After 30 min at room temperature, the plates were washed three times with wash buffer (0.05% Tween 20 in PBS). Serum samples were serially diluted in antibody diluent (1% BSA and 0.05% Tween 20 in PBS) and added to the washed plates (50µl/well). After incubation (RT, 2 h), the plates were washed three times and 1:10,000 diluted HRP-conjugated anti-mouse IgG antibody (catalog no. A28177; ThermoFisher) or anti-ferret IgG antibody (catalog no. ab112770; Abcam, USA) were added (100µl/well). After incubation (RT, 1h), the plates were washed six times followed by 30 min RT incubation with HRP substrate solution (100µl/well) prepared by adding a 10mg o-phenylenediamine dihydrochloride (OPD) tablet (catalog no. P8287; MilliporeSigma) to 20ml of phosphate-citrate buffer preparation (catalog no. P4922; MilliporeSigma). The reaction was stopped by adding 1 M sulfuric acid (100 µl/well), and the optical density was measured at 492 nm (OD492). Area under the curve (AUC) values were calculated using Prism8 software v.8.4.3 (GraphPad Software, USA). The baseline for AUC calculation was set as 0.1 to exclude non-specific signals from the AUC calculation. The OD492 of 0.1 is approximately 2 times the OD492 value from the control wells that were treated equally, but without the addition of diluted serum. Reciprocal dilutions (dilution factors) of the serum were used as x-values for the AUC calculation. Additionally, the level of secretory IgA in mice was also measured from bronchoalveolar lavage (BAL) fluid collected approximately three weeks-post boost immunization. The BAL fluids were serially diluted, and the level of IgA was measured as described above. HRP-conjugated anti-mouse IgA antibody (catalog no. ab97235; Abcam) was used.

#### Histopathology and immunohistochemistry

NBF-fixed mouse and ferret lungs were processed for histopathology and immunohistochemistry as previously described (*13*). Hematoxylin and eosin (H&E)-stained slides were examined from two mice or four ferrets per virus group at 5 days post-infection. Immunohistochemistry was done on the same sets of fixed tissues as the histopathology. For the mouse lung tissues, Influenza A virus and immune cell (neutrophil, B and T cells) distribution were measured by immunohistochemistry. A goat polyclonal primary anti-influenza A virus (catalog no. ab20841; Abcam, USA) was used to stain influenza NP proteins. Anti-CD19 antibody (catalog no. 90176S; Cell Signaling Technology, USA), anti-CD3 antibody (catalog no. ab16669; Abcam), and anti-Ly6G antibody (catalog no. ab210204; Abcam) were used to stain B cells, T cells, and neutrophils, respectively. For the ferret lung tissues, only Influenza A virus distribution was measured by immunohistochemistry. All slides were scanned on an Aperio ScanScope XT system (Aperio, USA), enabling whole-slide analysis.

#### Statistics

Prism8 software v.8.4.3 (GraphPad Software) was used for statistical evaluations of antibody levels and viral titers by ordinary ANOVA test. Tukey’s multiple comparison test was used as post hoc test to compared antibody levels between groups. Viral titers of the IM and IN groups were log-transformed and compared to Mock group using Dunnett’s multiple comparison test as post hoc test. Hierarchical clustering and additional analyses were performed using TIBCO Spotfire® Analyst 7.6.0 (TIBCO Software, Palo Alto, CA).

#### GMP manufacture and toxicology studies

The four vaccine virus (Supplementary Table 1) seed stocks were used for Good Manufacturing Practice (GMP) manufacturing of the vaccine components in certified Vero cells. A rabbit toxicology and immunogenicity study were performed under contract by Batelle. Thirty 4-to-7 month old New Zealand White Rabbits (2 to 4 kg) were employed for the study. A group of 10 animals received saline administered by IN (240 µL; 120 µL per naris) and IM (n = 10; 5 male & 5 female), another group of 10 received the GMP-manufactured vaccine IN (20ug of each antigen; 80ug total), and the final group of 10 received the GMP-manufactured vaccine IM (20ug of each antigen; 80ug total). Each group consists of 5 male and 5 female rabbits. First dose of vaccine was delivered on Day 1 and a boost was delivered on Day 29. Body weight measurements were made throughout the study. Body temperatures were measured prior to vaccination and at 6 and 24 hours following vaccination on days 1 and 29. Blood collection was performed before the study and on days 1, 8, 15, 30 and 45. Urine was collected prior to the study and on days 1, 8, 15, 30 and 45. Animals were sacrifice on day 45 (n = 30) for histopathological analyses from harvested tissues. Good Laboratory Practice (GLP) was followed for these experiments. Body weight and temperature were evaluated for any differences occurring during the study. Collected blood was evaluated for clinical chemistry and immunogenicity. Urinalysis was performed on collected urine samples over the course of the study. Harvested tissues underwent histopathologic evaluation for signs of toxicity.

---

**Fig. S1:**
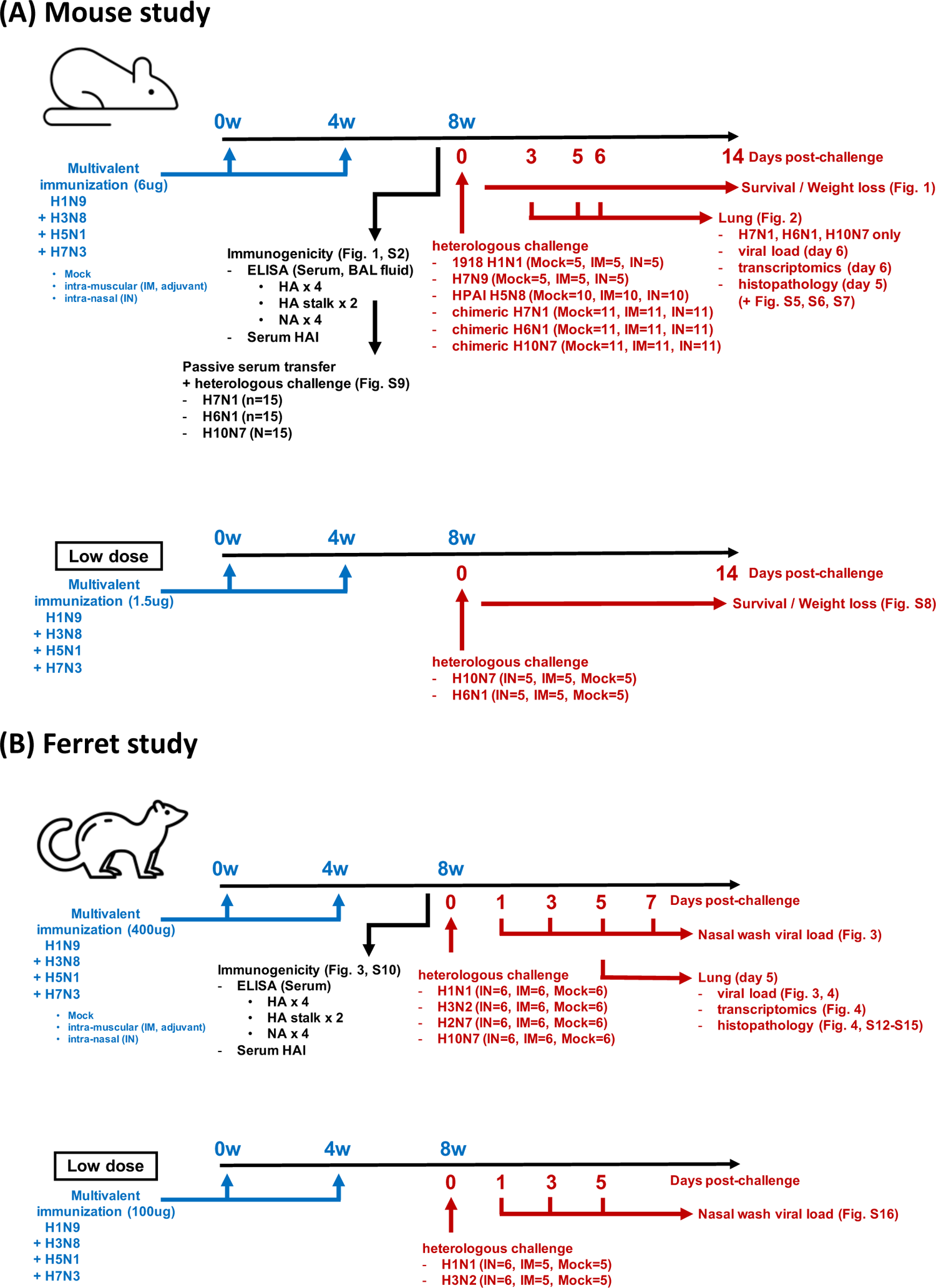
Animal study description

**Fig. S2:**
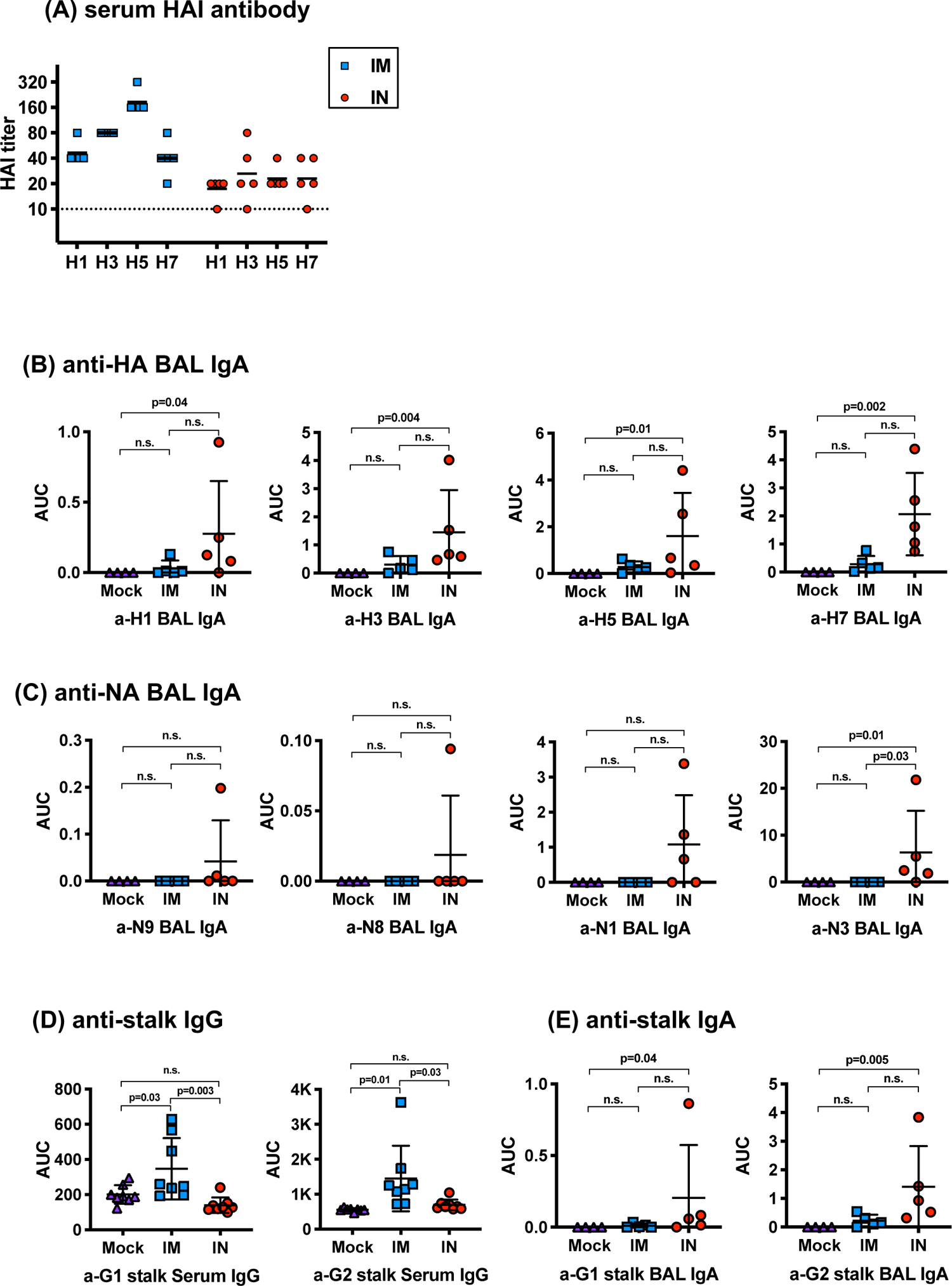
Immunogenicity in mice. Balb/c mice were intramuscularly (IM) or intranasally (IN) immunized twice with the BPL-inactivated vaccine. Serum and bronchoalveolar lavage fluid (BALF) samples were collected 3-weeks-post the second immunization. **(A)** Hemagglutination inhibition (HAI) antibody titers against the four vaccine antigens were measured from serum samples from IM- or IN-vaccinated mice. The dashed line shows the detection limit of the HAI assay used. IgA levels against **(B)** four vaccine hemagglutinin (HA) antigens and **(C)** four vaccine neuraminidase (NA) antigens were measured in mock-, IM-, or IN-vaccinated mice using ELISA. **(D)** Serum IgG or **(E)** BALF IgA antibody levels against group-1 and group-2 HA stalk were measured using ELISA. Error bars represent standard deviation. Kruskal-Wallis test and post hoc Dunn’s multiple comparison test were used to compare BALF IgA levels between groups. Ordinary ANOVA test and post hoc Tukey’s multiple comparison test were used to compared serum IgG levels between groups. **n.s**; not significant, **AUC**; area under the curve

**Supplementary Fig. S3:**
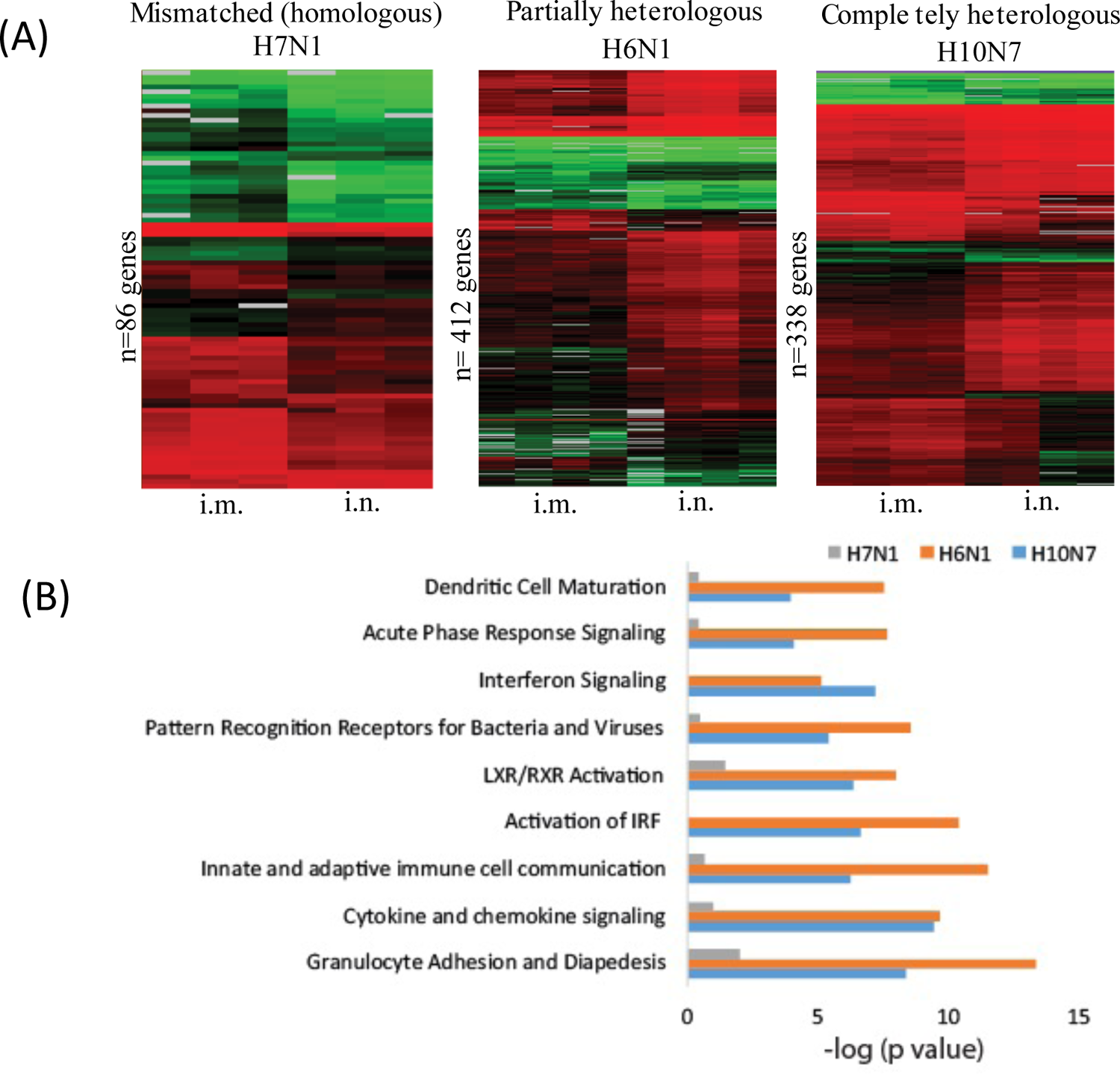
Differential host response to influenza infection in IN and IM vaccinated mice. (A) Heatmap represents transcripts showing differential expression (≥2-fold difference in median expression level, p<0.05) between IN and IM vaccinated animals challenged with either H7N1, H6N1 or H10N7 influenza virus on day 6 post-infection. Genes with increased expression are show in red, genes with no change as black, and genes showing decreased expression in green. **(B)** Gene enrichment analysis of genes showing differential expression between IN and IM vaccinated animals.

**Supplementary Fig. S4:**
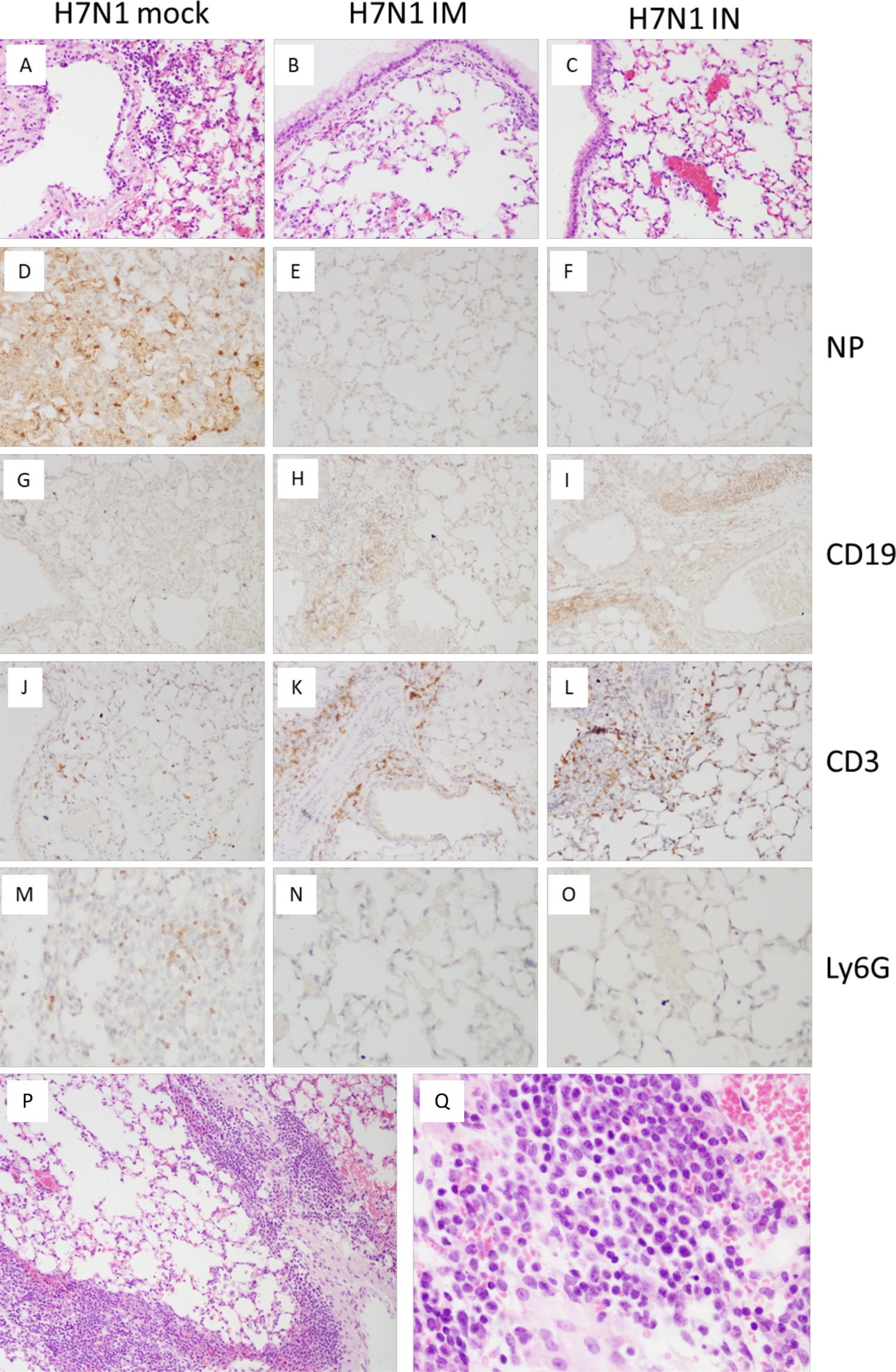
Mouse H7N1 pathology. **(A-C)**. H&E stains of lung at day 5 post challenge. Mock-vaccinated mice (**A**) challenged with the H7N1 virus showed an extensive primary viral pneumonia, affecting >50% of lung with necrotizing bronchitis and bronchiolitis, and alveolitis with a neutrophil-rich inflammatory infiltrate, while IM-vaccinated mice (**B**) and IN-vaccinated mice (**C**) showed no evidence of pneumonia or inflammatory infiltrates. Original magnifications 40x. (**D-F**). Mock-vaccinated mice (**D**) challenged with the H7N1 virus showed extensive viral antigen (nucleoprotein, NP) staining in alveolar epithelial cells and alveolar macrophages, while IM-vaccinated mice (**E**) and IN-vaccinated mice (**F**) showed no viral antigen staining. Original magnifications 100x. (**G-I**). Immunostaining for CD19+ B cells. Mock-vaccinated mice (**G**) challenged with the H7N1 virus showed few, scattered CD19+ cells, while IM-vaccinated mice (**H**) and IN-vaccinated mice (**I**) showed prominent perivascular and peribronchiolar aggregates of CD19+ cells. Original magnifications 40x. (**J-L**). Immunostaining for CD3+ T cells. Mock-vaccinated mice (**J**) challenged with the H7N1 virus showed few, scattered CD3+ cells, while IM-vaccinated mice (**K**) and IN-vaccinated mice (**L**) showed prominent perivascular and peribronchiolar aggregates of CD3+ cells. Original magnifications 40x. (**M-O**). Immunostaining for neutrophils with the Ly6G antibody. Mock-vaccinated mice (**M**) challenged with the H7N1 virus showed many Ly6G+ neutrophils within the lung parenchyma, while IM-vaccinated mice (**N**) and IN-vaccinated mice (**O**) showed no neutrophils. Original magnifications 100x. (**P-Q**). Large aggregates of plasma cells (staining negatively for CD19) were observed in the lungs of vaccinated animals but not in lungs of mock-vaccinated animals. Original magnifications 40 x (**P**) and 200x (**Q**).

**Supplementary Fig. S5:**
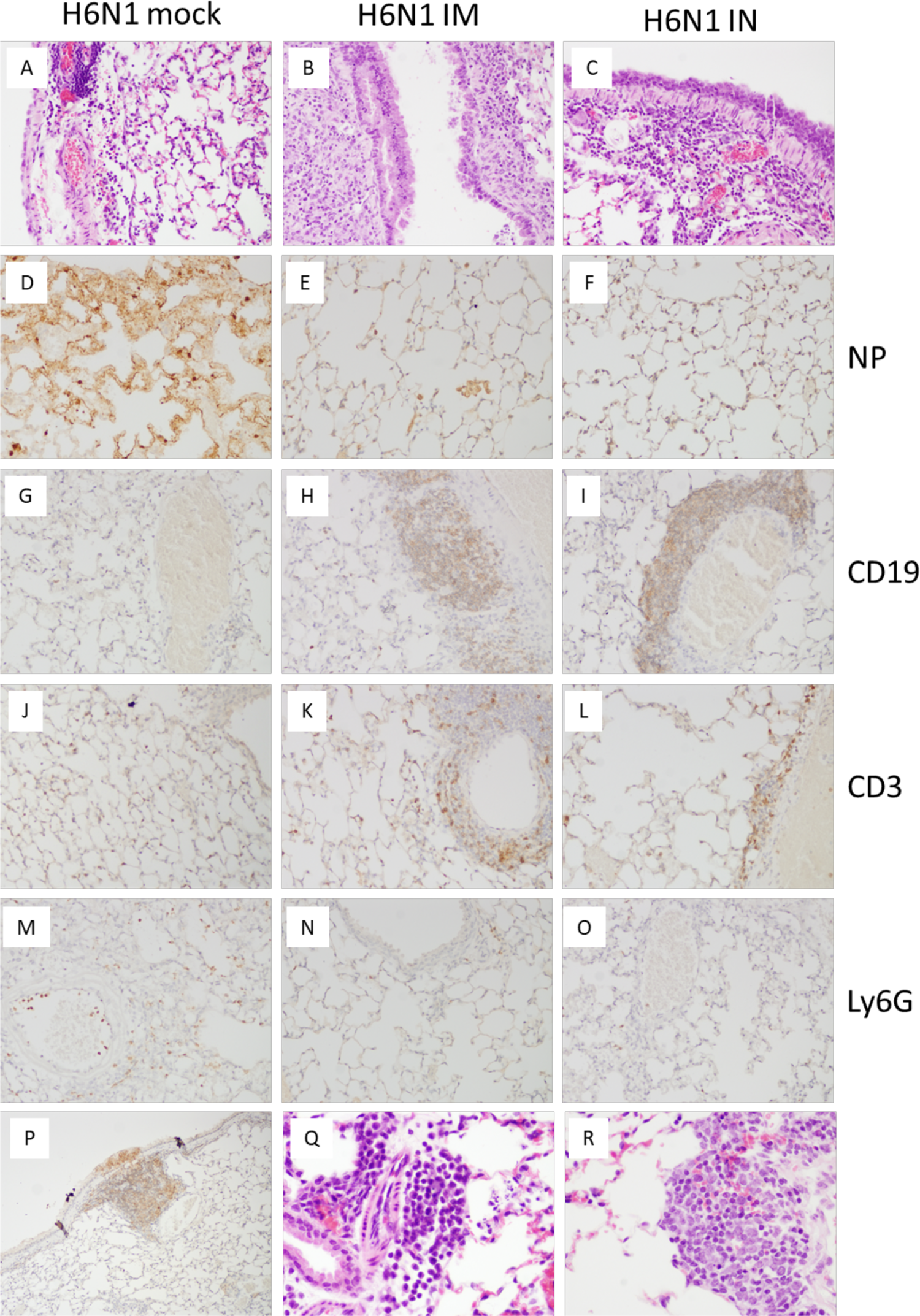
Mouse H6N1 pathology. **(A-C)**. H&E stains of lung at day 5 post challenge. Mock-vaccinated mice (**A**) challenged with the H6N1 virus showed an extensive primary viral pneumonia, affecting >50% of lung with necrotizing bronchitis and bronchiolitis, and alveolitis with a neutrophil-rich inflammatory infiltrate, while IM-vaccinated mice (**B**) and IN-vaccinated mice (**C**) showed little evidence of pneumonia or inflammatory infiltrates. The respiratory epithelium of vaccinated mice showed extensive reproliferation, and submucosal lymphoid aggregates were prominent. Original magnifications 40x. (**D-F**). Mock-vaccinated mice (**D**) challenged with the H7N1 virus showed extensive viral antigen (nucleoprotein, NP) staining in alveolar epithelial cells and alveolar macrophages, while IM-vaccinated mice (**E**) and IN-vaccinated mice (**F**) showed little viral antigen staining, mostly in alveolar macrophages. Original magnifications 100x. (**G-I**). Immunostaining for CD19+ B cells. Mock-vaccinated mice (**G**) challenged with the H7N1 virus showed few, scattered CD19+ cells, while IM-vaccinated mice (**H**) and IN-vaccinated mice (**I**) showed prominent perivascular and peribronchiolar aggregates of CD19+ cells. Original magnifications 40x. (**J-L**). Immunostaining for CD3+ T cells. Mock-vaccinated mice (**J**) challenged with the H7N1 virus showed few, scattered CD3+ cells, while IM-vaccinated mice (**K**) and IN-vaccinated mice (**L**) showed prominent perivascular and peribronchiolar aggregates of CD3+ cells. Original magnifications 40x. (**M-O**). Immunostaining for neutrophils with the Ly6G antibody. Mock-vaccinated mice (**M**) challenged with the H7N1 virus showed many Ly6G+ neutrophils within the lung parenchyma and margination from small blood vessels, while IM-vaccinated mice (**N**) and IN-vaccinated mice (**O**) showed very few neutrophils. Original magnifications 100x. (**P-R**). IM-vaccinated animals showed small foci of intra-epithelial CD19+ B cells (**P**). Aggregates of plasma cells (staining negatively for CD19) were observed in the lungs of vaccinated animals but not in lungs of mock-vaccinated animals (**Q-R**). Original magnifications 40 x (**P**) and 200x (**Q-R**).

**Supplementary Fig. S6:**
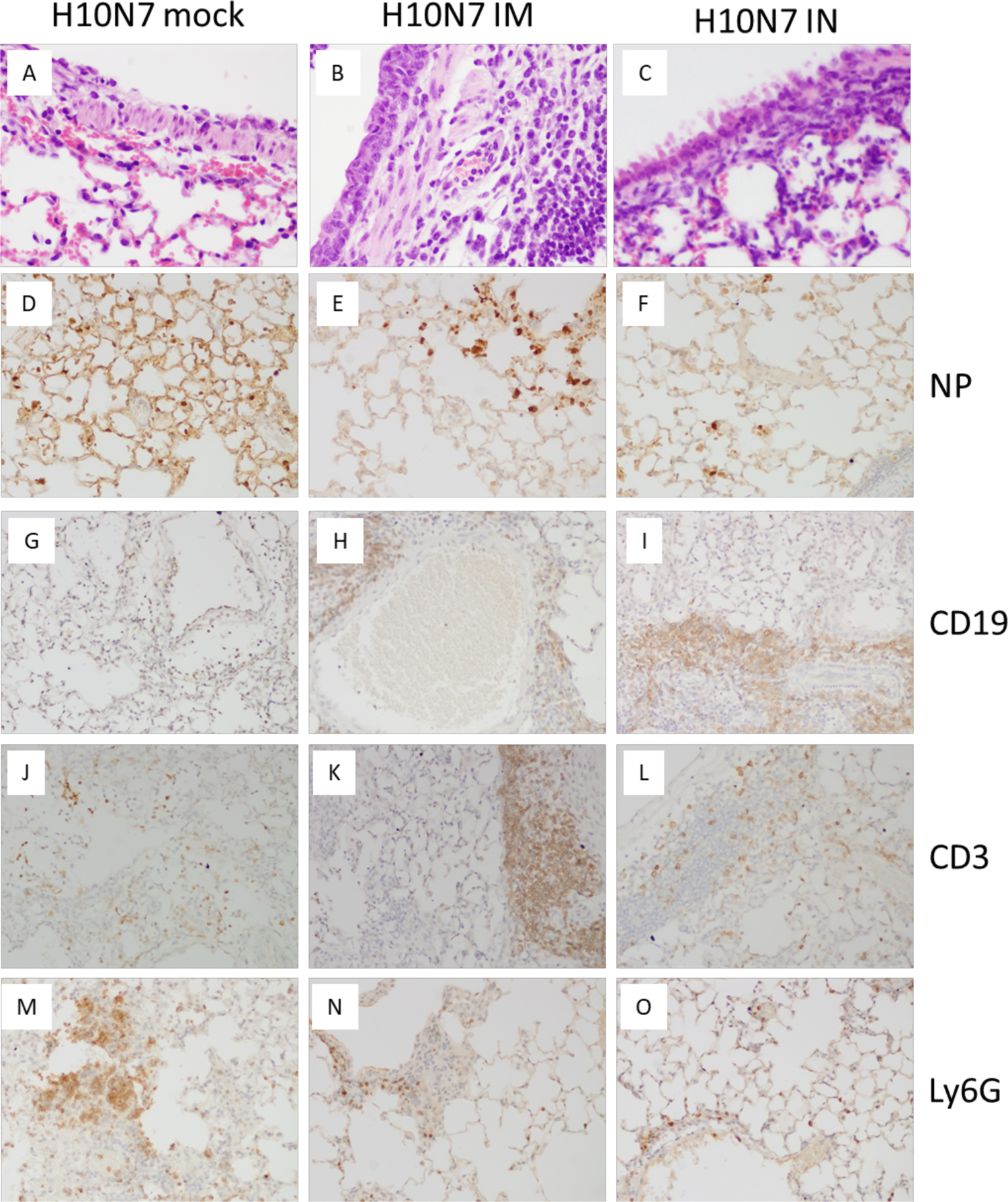
Mouse H10N7 pathology. **(A-C)**. H&E stains of lung at day 5 post challenge. Mock-vaccinated mice (**A**) challenged with the H10N7 virus showed an extensive primary viral pneumonia, affecting >50% of lung with necrotizing bronchitis and bronchiolitis, and alveolitis with a neutrophil-rich inflammatory infiltrate, while IM-vaccinated mice (**B**) and IN-vaccinated mice (**C**) showed little evidence of pneumonia or inflammatory infiltrates. The respiratory epithelium of vaccinated mice showed extensive reproliferation with prominent mitotic figures (**B**), and submucosal lymphoid aggregates were prominent. Original magnifications 40x. (**D-F**). Mock-vaccinated mice (**D**) challenged with the H7N1 virus showed extensive viral antigen (nucleoprotein, NP) staining in alveolar epithelial cells and alveolar macrophages, while IM-vaccinated mice (**E**) and IN-vaccinated mice (**F**) showed some viral antigen staining, mostly in alveolar macrophages. Original magnifications 100x. (**G-I**). Immunostaining for CD19+ B cells. Mock-vaccinated mice (**G**) challenged with the H7N1 virus showed few, scattered CD19+ cells, while IM-vaccinated mice (**H**) and IN-vaccinated mice (**I**) showed prominent perivascular and peribronchiolar aggregates of CD19+ cells. Original magnifications 40x. (**J-L**). Immunostaining for CD3+ T cells. Mock-vaccinated mice (**J**) challenged with the H7N1 virus showed few, scattered CD3+ cells, while IM-vaccinated mice (**K**) and IN-vaccinated mice (**L**) showed prominent perivascular and peribronchiolar aggregates of CD3+ cells. Original magnifications 40x. (**M-O**). Immunostaining for neutrophils with the Ly6G antibody. Mock-vaccinated mice (M) challenged with the H7N1 virus showed large infiltrates Ly6G+ neutrophils within the lung parenchyma and margination from small blood vessels, while IM-vaccinated mice (**N**) and IN-vaccinated mice (**O**) showed only few neutrophils, mostly in peribronchiolar locations. Original magnifications 100x.

**Supplementary Fig. S7:**
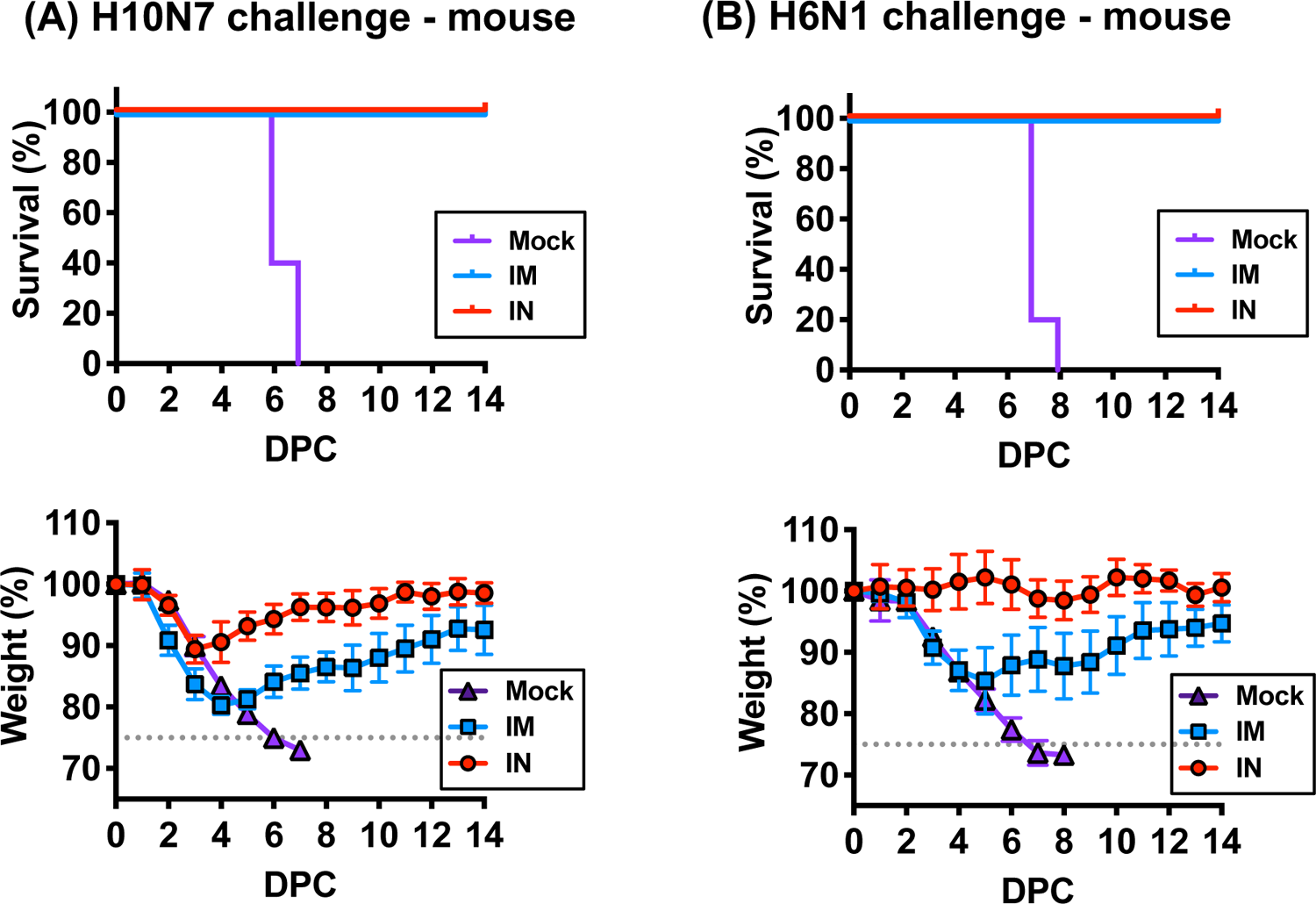
Protection with low dose (¼) immunization in mice. Balb/c mice were intramuscularly (IM) or intranasally (IN) immunized twice with the BPL-inactivated vaccine at ¼ dose (i.e. 1.5ug total) followed by 10x mouse 50% lethal dose (LD50) challenge. Percent survival and percent weight loss in each group (n=5) following **(A)** H10N7 or **(B)** H6N1 virus challenge are shown. Error bars represent standard deviation. **DPC**; days post-challenge

**Supplementary Fig. S8:**
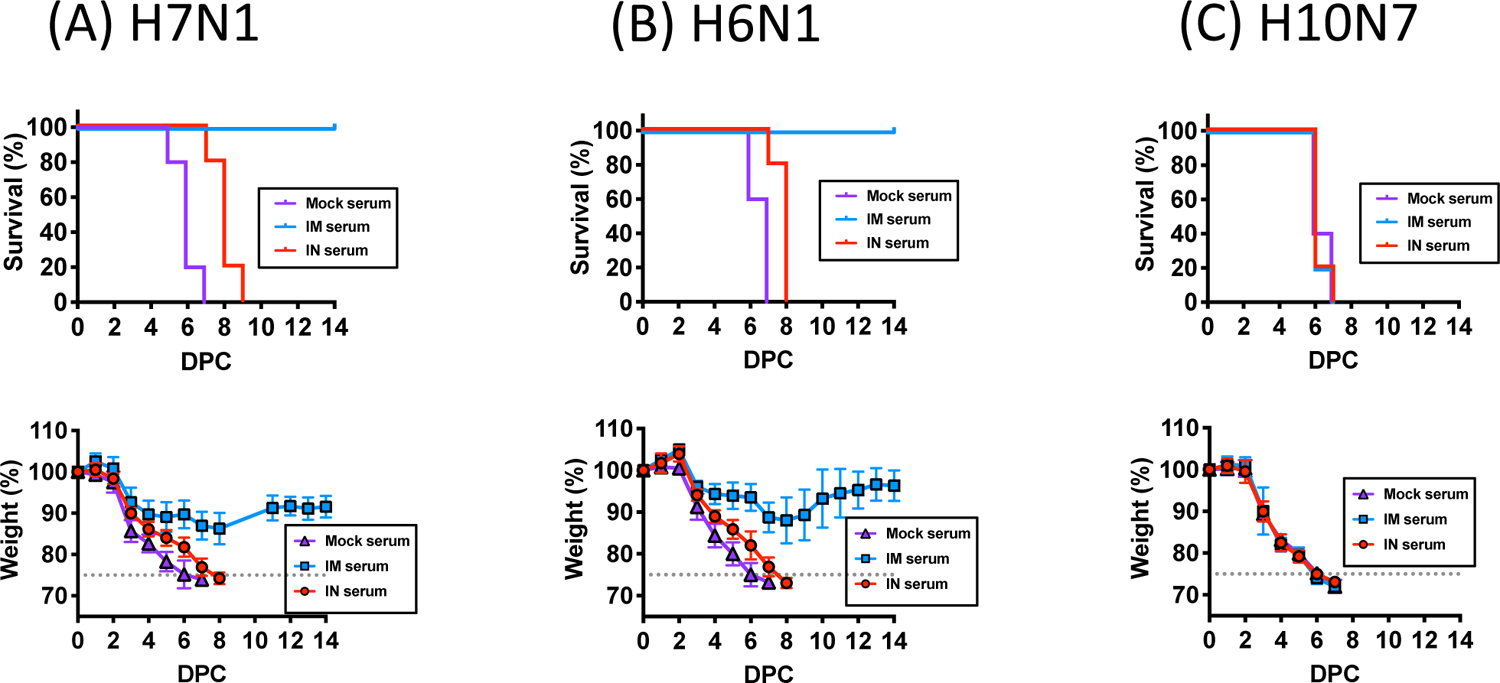
Protection by passive serum transfer in mice. Serum from mock-, IM-, or IN-vaccinated mice was injected intraperitoneally (200ul per animal) 1 day prior to 10x mouse 50% lethal dose (LD50) challenge. Percent survival and percent weight loss in each group following **(A)** H7N1, **(B)** H6N1, and **(C)** H10N7 virus challenge are shown. Error bars represent standard deviation. **DPC**; days post-challenge

**Supplementary Fig. S9:**
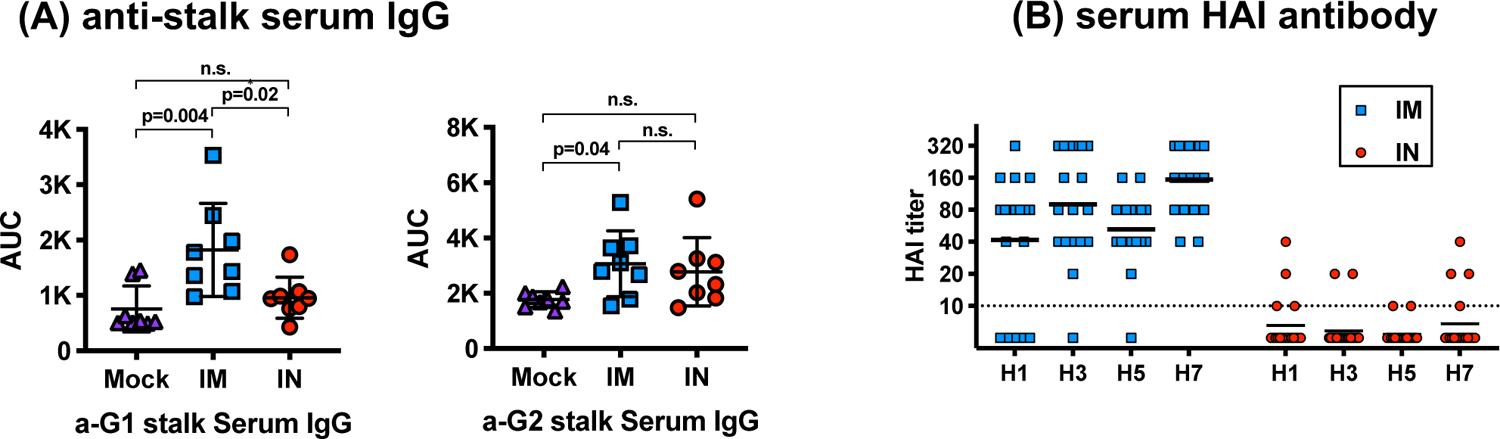
Immunogenicity in ferrets. Ferrets were intramuscularly (IM) or intranasally (IN) immunized twice with the BPL-inactivated vaccine. Serum samples were collected 3-weeks-post the second immunization. **(A)** Serum IgG levels against group-1 and group-2 HA stalk were measured by ELISA. Error bars represent standard deviation. Ordinary ANOVA test and post hoc Tukey’s multiple comparison test were used to compared serum IgG levels between groups. **(B)** Hemagglutination inhibition (HAI) antibody titers against vaccine antigens were measured IM- or IN-vaccinated ferrets. The dashed line shows the detection limit of the HAI assay used. **n.s**; not significant, **AUC**; area under the curve

**Supplementary Fig. S10:**
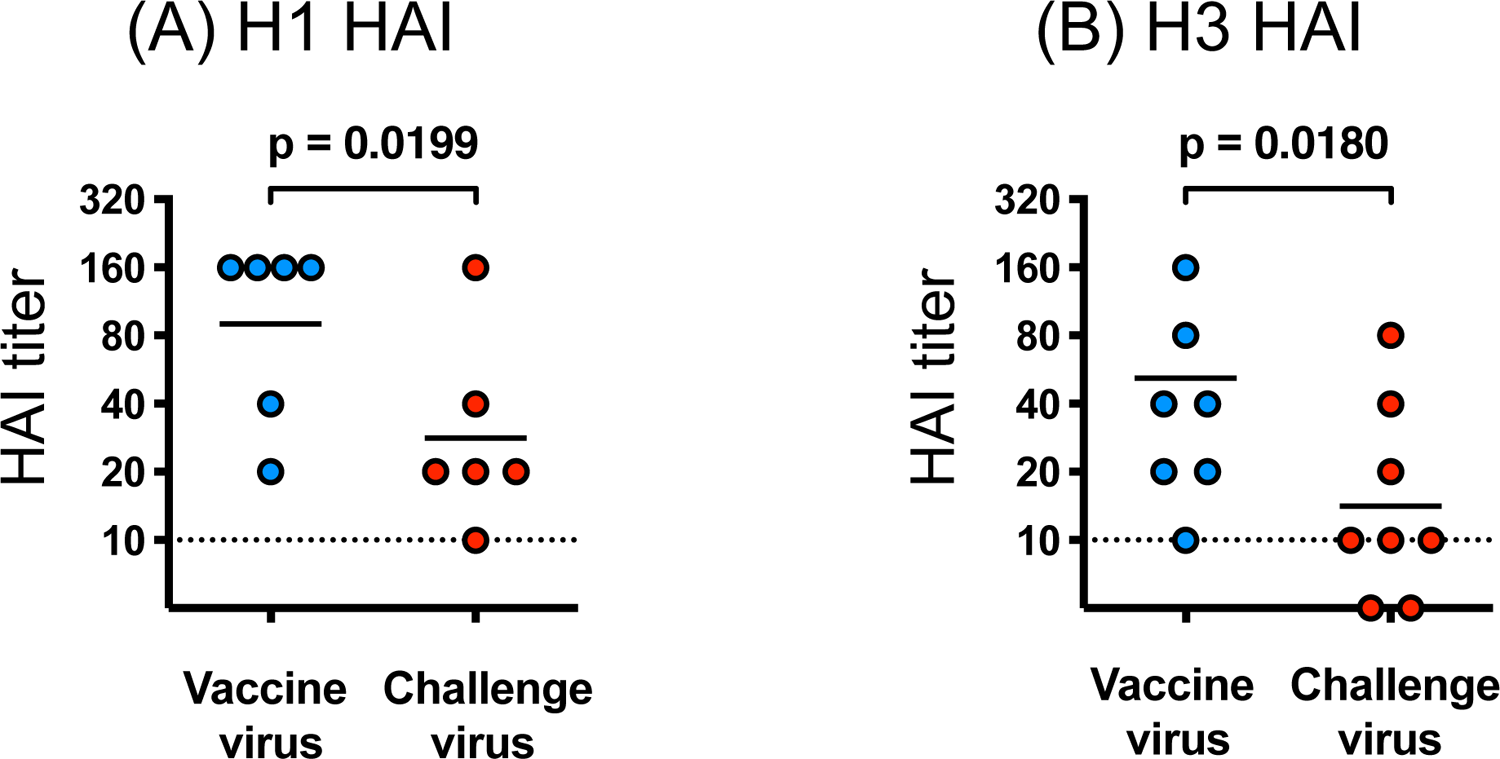
Antigenic differences between the vaccine strain and the challenge strain measured by hemagglutination inhibition (HAI) assays. Ferrets were intramuscularly (IM) or intranasally (IN) immunized twice with the BPL-inactivated vaccine. Serum samples were collected 3-weeks-post the second immunization. **(A)** Serum HAI titers were measured against the vaccine strain A/mallard/Ohio/265/1987 (H1N9) and the challenge strain A/swine/Iowa/1931 (H1N1). **(B)** Serum HAI titers were measured against the vaccine strain A/pintail/Ohio/339/1987 (H3N8) and the challenge strain A/Port Chalmers/1973 (H3N2). A two-tailed paired t-test was used on log-transformed HAI titers for comparison. Bars represent geometric means.

**Supplementary Fig. S11:**
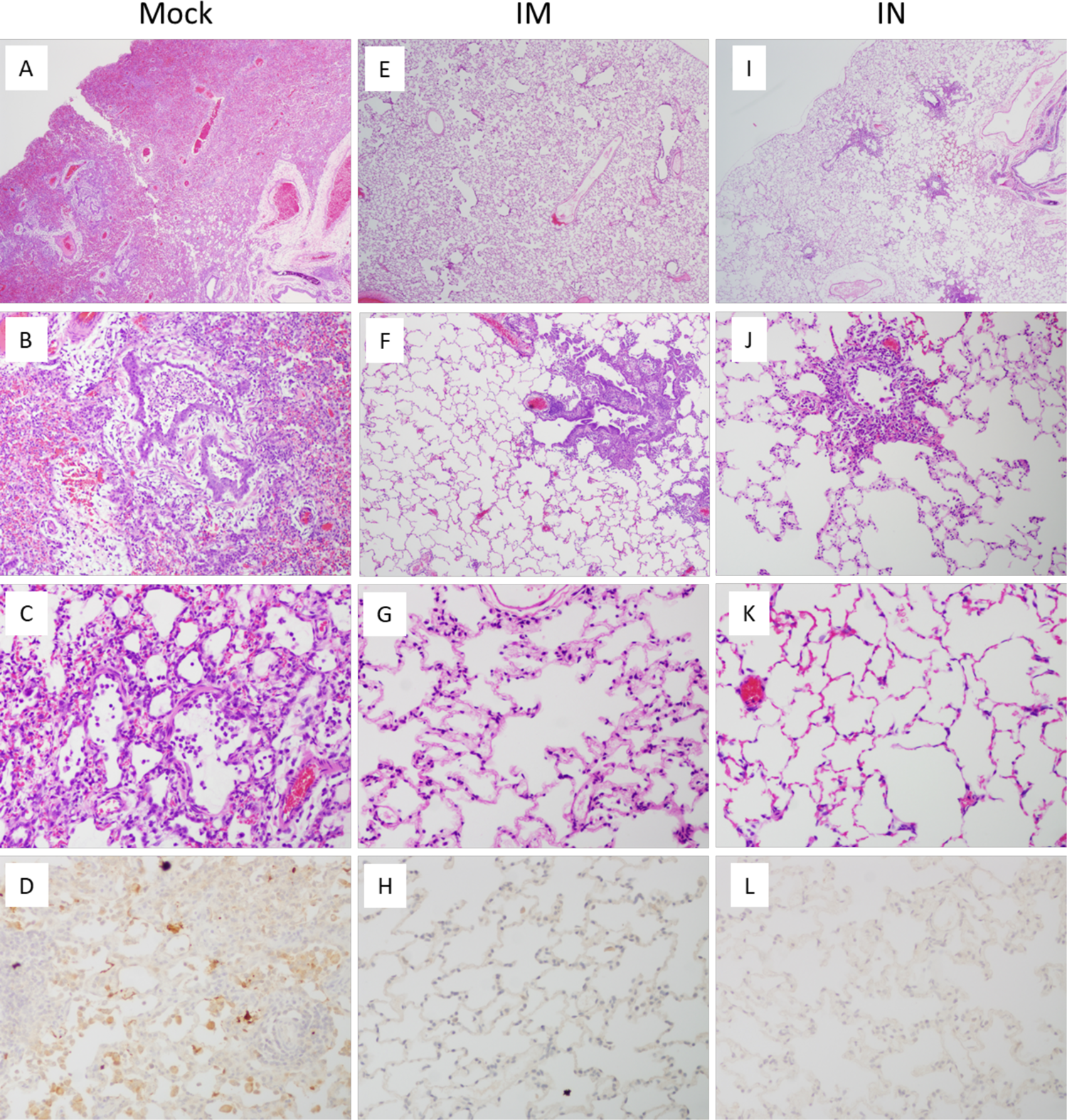
Ferret H1N1 pathology. **(A-C)**. H&E stains of lung at day 5 post challenge. Mock-vaccinated ferrets (**A-D**) challenged with the swine/1931 H1N1 virus showed an extensive primary viral pneumonia (**A**), affecting >50% of lung with necrotizing bronchitis and bronchiolitis (**B**), and alveolitis with a neutrophil-rich inflammatory infiltrate (**C**). Immunostaining revealed extensive viral antigen (nucleoprotein, NP) in alveolar epithelial cells and alveolar macrophages (**D**). IM-vaccinated ferrets (**E-H**) challenged with the swine/1931 H1N1 virus showed no pneumonia (**E**), with multifocal, mild bronchiolitis (**F**), no alveolitis (**G**), and no viral antigen staining (**H**). IN-vaccinated ferrets (**I-L**) challenged with the swine/1931 H1N1 virus showed no pneumonia (**I**), with multifocal, mild bronchiolitis (**J**), no alveolitis (**K**), and no viral antigen staining (**L**). Original magnifications: 40x (**A, B, E, F, I, J**), and 100x (**C, D, G, H, K, L**).

**Supplementary Fig. S12:**
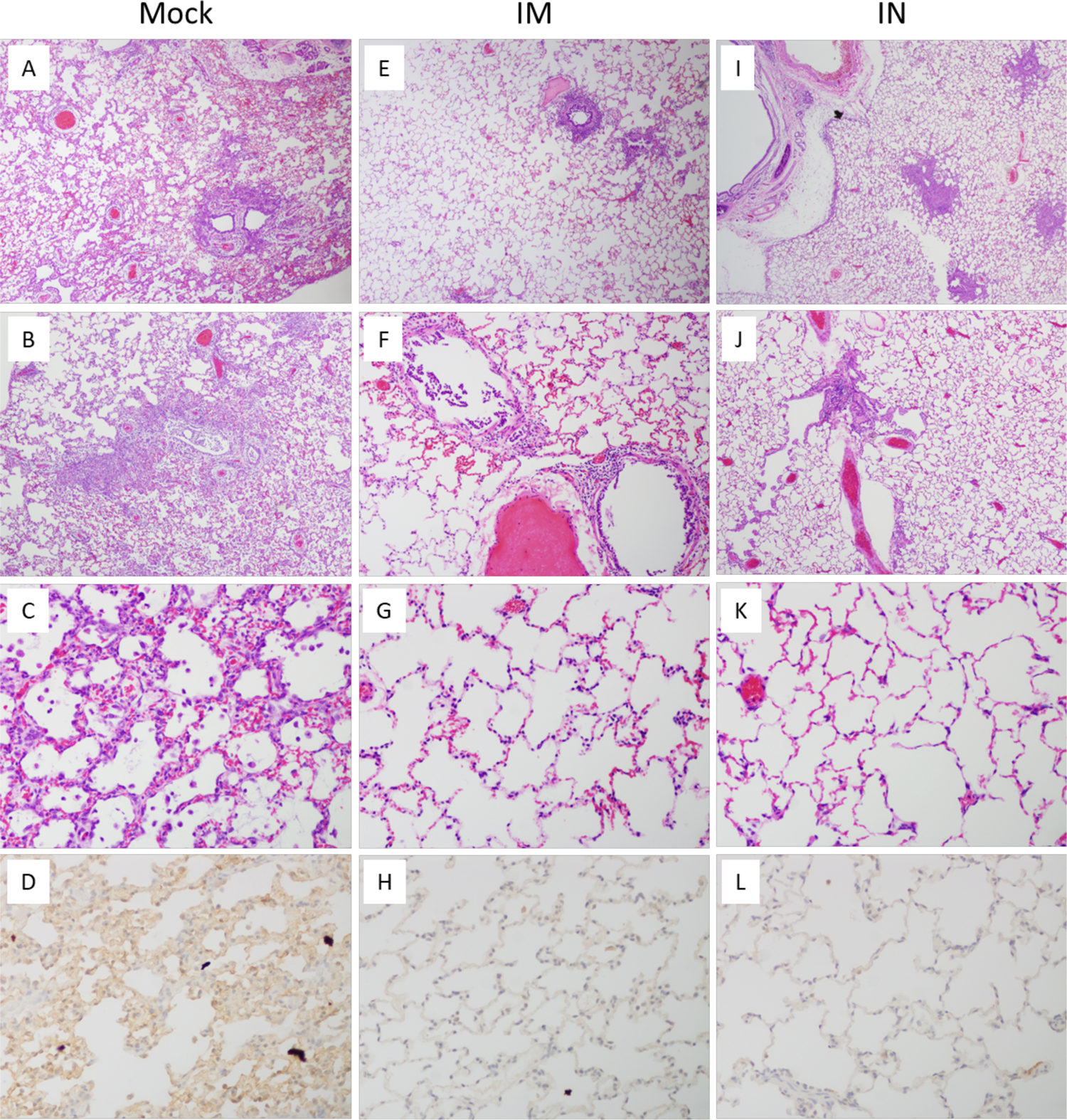
Ferret H3N2 pathology. **(A-C)**. H&E stains of lung at day 5 post challenge. Mock-vaccinated ferrets (**A-D**) challenged with the human A/Port Chalmers/1973 H3N2 virus showed an extensive primary viral pneumonia (**A**), affecting >50% of lung with necrotizing bronchitis and bronchiolitis (**B**), and alveolitis with a neutrophil-rich inflammatory infiltrate (**C**). Immunostaining revealed extensive viral antigen (nucleoprotein, NP) in alveolar epithelial cells and alveolar macrophages (**D**). IM-vaccinated ferrets (**E-H**) challenged with the H3N2 virus showed no pneumonia (**E**), with multifocal, mild bronchiolitis (**F**), no alveolitis (**G**), and no viral antigen staining (**H**). IN-vaccinated ferrets (**I-L**) challenged with the H3N2 virus showed no pneumonia (**I**), with multifocal, mild bronchiolitis and submucosal lymphoid aggregates (**J**), no alveolitis (**K**), and no viral antigen staining (**L**). Original magnifications: 40x (**A, B, E, F, I, J**), and 100x (**C, D, G, H, K, L**).

**Supplementary Fig. S13:**
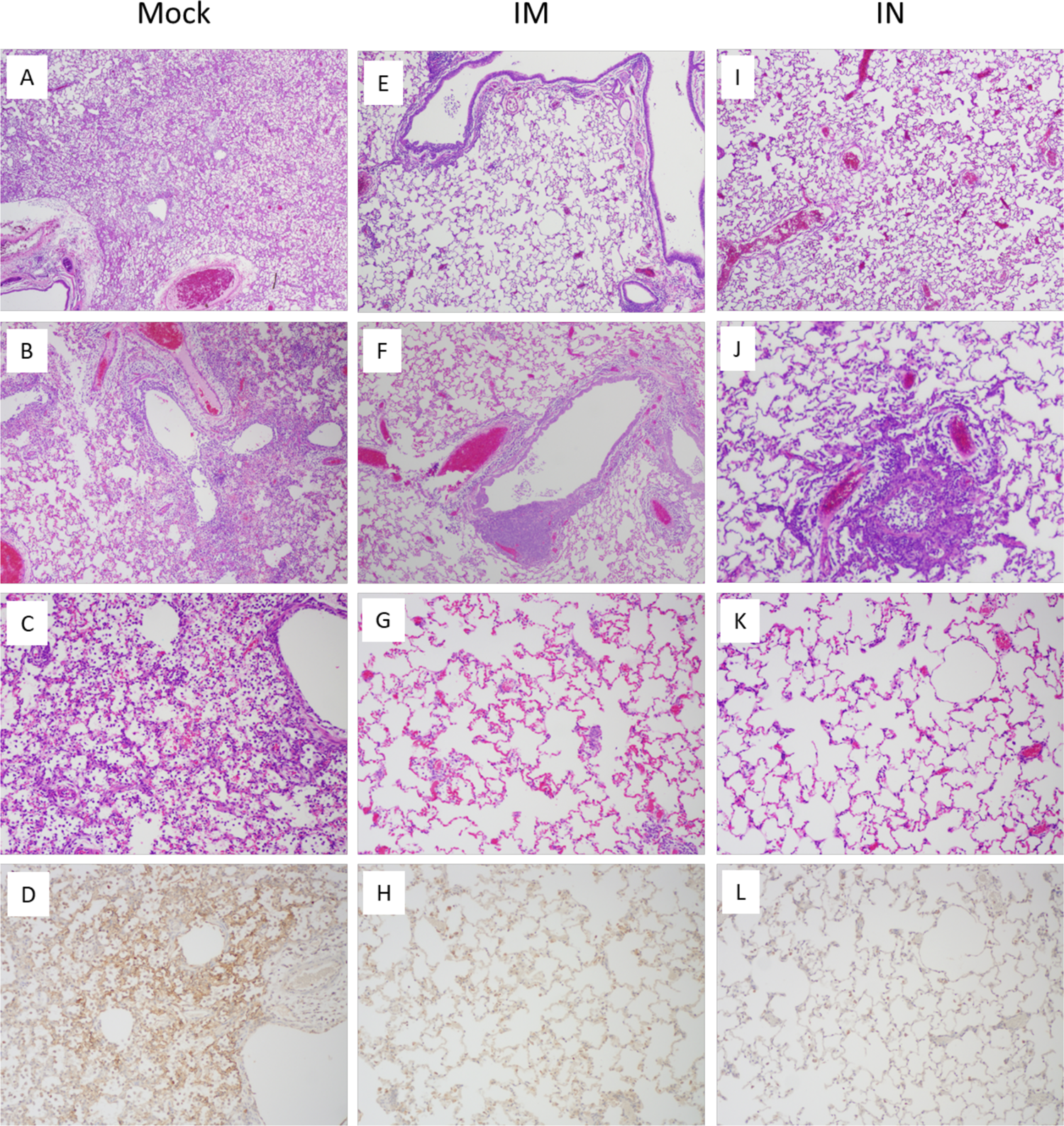
Ferret H2N7 pathology. **(A-C)**. H&E stains of lung at day 5 post challenge. Mock-vaccinated ferrets (**A-D**) challenged with the chimeric H2N7 virus showed an extensive primary viral pneumonia (**A**), affecting >50% of lung with necrotizing bronchitis and bronchiolitis (**B**), and alveolitis with a neutrophil-rich inflammatory infiltrate (**C**). Immunostaining revealed extensive viral antigen (nucleoprotein, NP) in alveolar epithelial cells and alveolar macrophages (**D**). IM-vaccinated ferrets (**E-H**) challenged with the H3N2 virus showed no pneumonia (**E**), with multifocal, mild bronchiolitis with prominent submucosal lymphoid aggregates (**F**), no alveolitis (**G**), and no viral antigen staining (**H**). IN-vaccinated ferrets (**I-L**) challenged with the H3N2 virus showed no pneumonia (**I**), with multifocal, mild bronchiolitis and submucosal lymphoid aggregates (**J**), no alveolitis (**K**), and no viral antigen staining (**L**). Original magnifications: 40x (**A, B, E, F, I, J**), and 100x (**C, D, G, H, K, L**).

**Supplementary Fig. S14:**
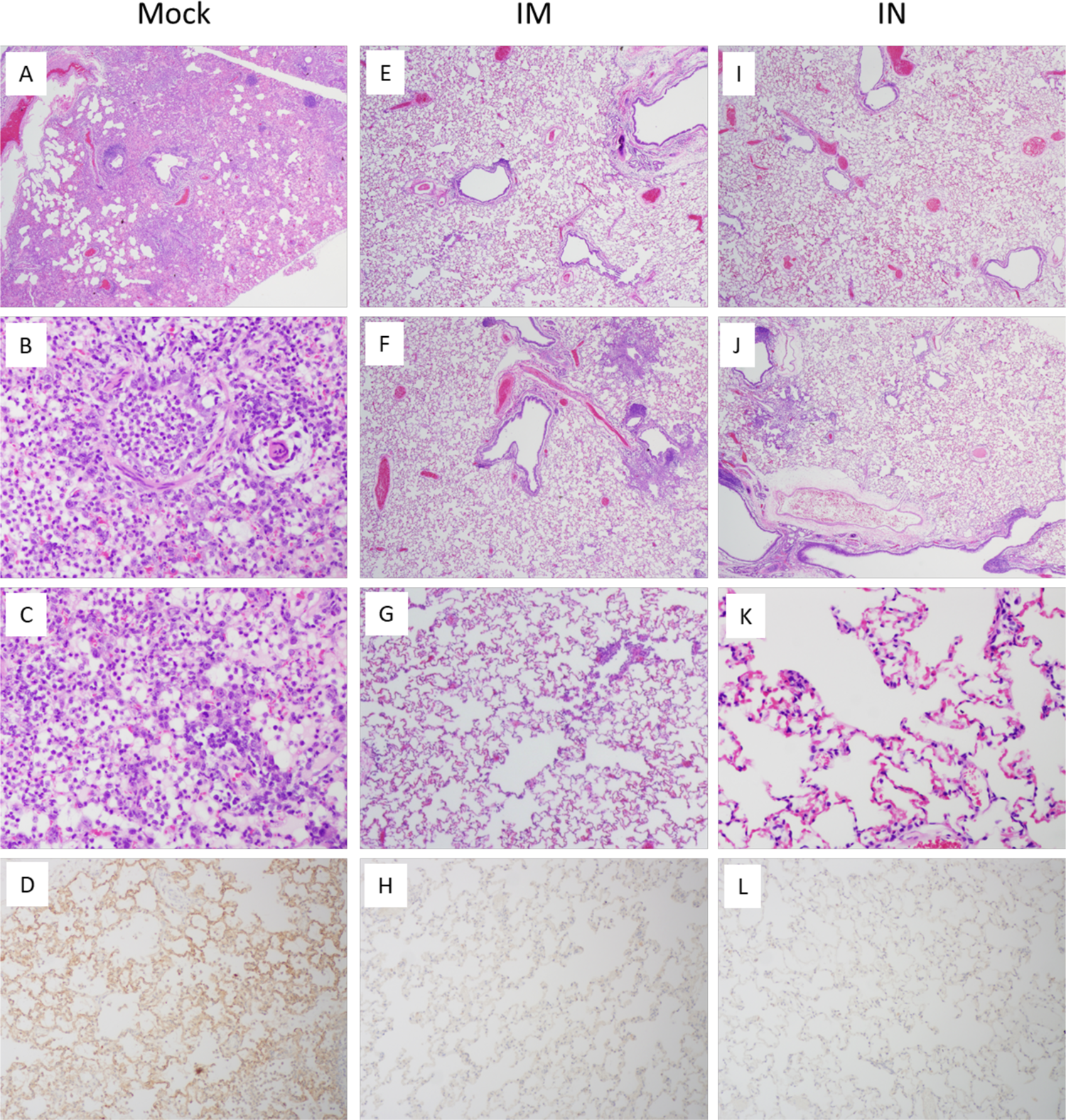
Ferret H10N7 pathology. **(A-C)**. H&E stains of lung at day 5 post challenge. Mock-vaccinated ferrets (**A-D**) challenged with the chimeric H10N7 virus showed an extensive primary viral pneumonia (**A**), affecting >50% of lung with necrotizing bronchitis and bronchiolitis (**B**), and alveolitis with a neutrophil-rich inflammatory infiltrate (**C**). Immunostaining revealed extensive viral antigen (nucleoprotein, NP) in alveolar epithelial cells and alveolar macrophages (**D**). IM-vaccinated ferrets (**E-H**) challenged with the H3N2 virus showed no pneumonia (**E**), with multifocal, mild bronchiolitis with prominent submucosal lymphoid aggregates (**F**), no alveolitis (**G**), and no viral antigen staining (**H**). IN-vaccinated ferrets (**I-L**) challenged with the H3N2 virus showed no pneumonia (**I**), with multifocal, mild bronchiolitis and submucosal lymphoid aggregates (**J**), no alveolitis (**K**), and no viral antigen staining (**L**). Original magnifications: 40x (**A, B, E, F, I, J**), and 100x (**C, D, G, H, K, L**).

**Supplementary Fig. S15:**
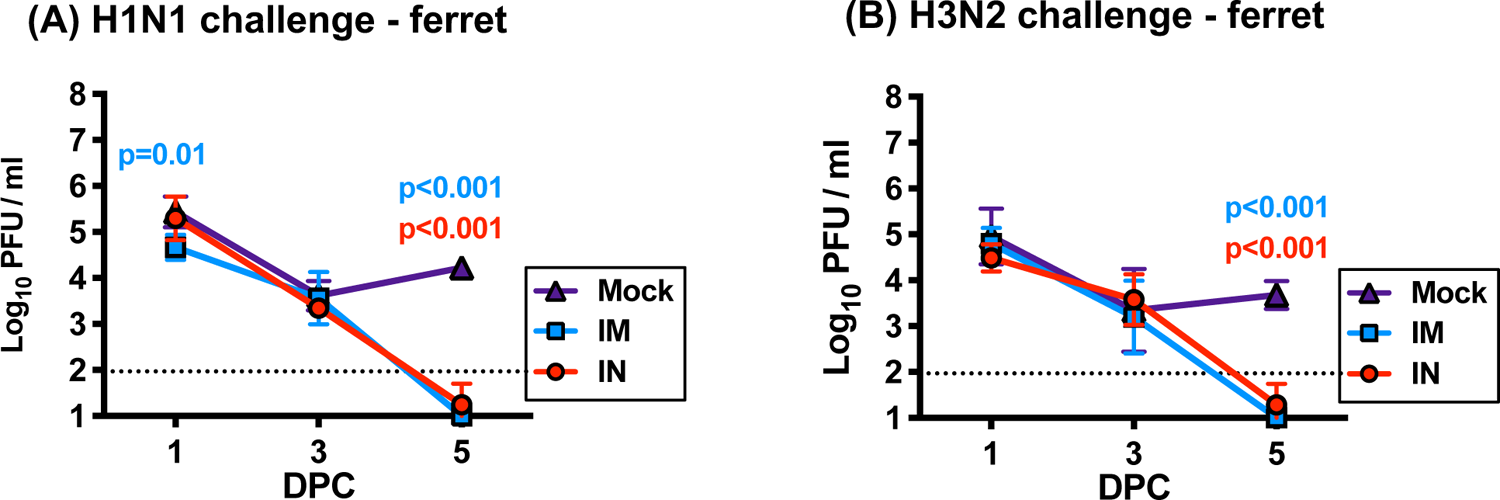
Protection with low dose (¼) immunization in ferrets. Ferrets were intramuscularly (IM) or intranasally (IN) immunized twice with the BPL-inactivated vaccine at ¼ dose (i.e.100ug total) and challenged with **(A)** A/swine/Iowa/1931 (H1N1) virus or **(B)** A/Port Chalmers/1973 (H3N2) virus. Nasal wash samples were collected at days 1, 3, and 5, and the titer was measured using plaque assay. Ordinary ANOVA test and post hoc Dunnett’s multiple comparison test were used to compare viral titers in IM- or IN-vaccinated ferrets to mock-vaccinated ferrets. Error bars represent geometric standard deviation. Dashed lines show the detection limit of the plaque assay used. **DPC**; days post-challenge

**Supplementary Fig. S16:**
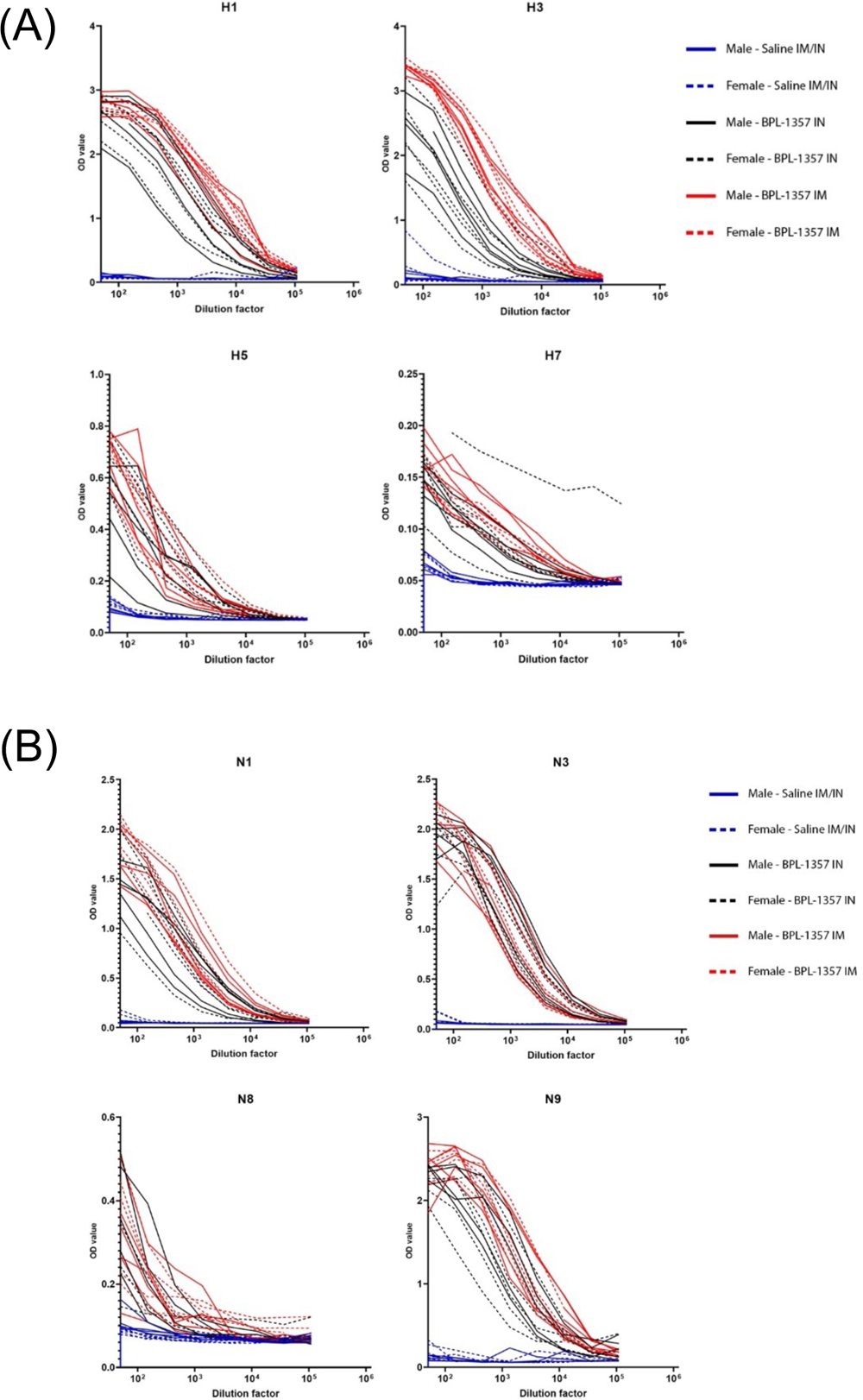
Immunogenicity of the vaccine in New Zealand white rabbits. Good Manufacturing Practice (GMP) manufacturing of the multivalent BPL-inactivated vaccine was performed in certified Vero cells. Immunogenicity of the GMP-manufactured vaccine was tested in thirty New Zealand White Rabbits following two immunizations 4 weeks apart. Ten rabbits were intranasally (IN) immunized with the GMP vaccine (n=10; 5 male, 5 female). Ten rabbits were intramuscularly (IM) immunized with the GMP vaccine (n=10; 5 male, 5 female). Ten control rabbits received saline administered by IN (240 µL; 120 µL per naris) and IM (n = 10; 5 male & 5 female). Serum samples were collected 16-days-post the second immunization, and the serum IgG levels against **(A)** four vaccine hemagglutinin (HA) antigens and **(B)** four vaccine neuraminidase (NA) antigens were measured using ELISA.

**Supplementary Table S1.**
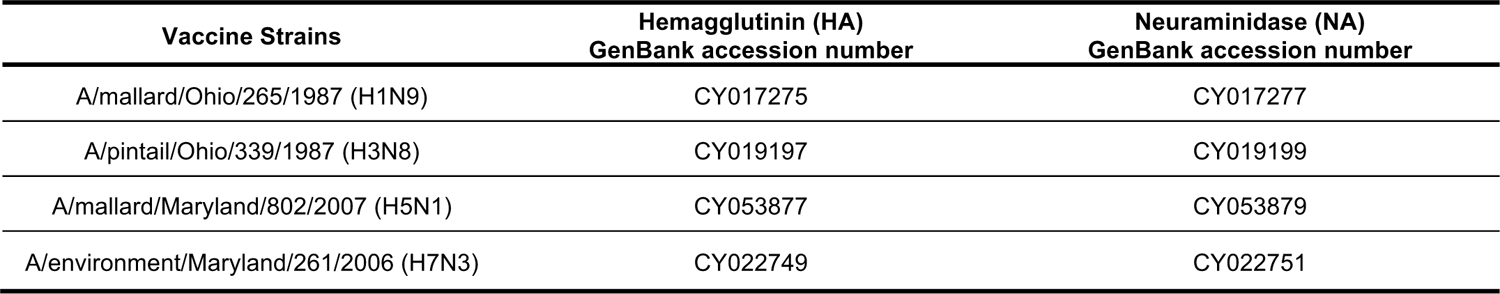
Vaccine strains

**Supplementary Table S2.** Challenge virus HA and NA identity to vaccine virus components

| **Mouse Challenge Influenza A viruses (10xLD<sub>50</sub>) and Identity to Vaccine Viruses** |  |  |
| --- | --- | --- |
| Subtype (strain) | Percent nucleotide (and amino acid) identities |  |
|  | HA | NA |
| H1N1 (A/Brevig Mission/1/1918) | 79.1% (92.8%) | 85.5% (91.5%) |
| H5N8 (A/chicken/Netherlands/EMC-3/2014) | 79.7% (85.6%) | 76.8% (84.8%) |
| H6N1 (recombinant Avian)<sup>§</sup> | 51.0% (63.1%) [vs H1]* | 92.1% (96.2%) |
| H7N1 (recombinant Avian)<sup>§</sup> | 100.0% (100%) | 92.1% (96.2) |
| H7N9 (A/Shanghai/1/2013) | 76.9% (84.8%) | 91.0% (95.5%) |
| H10N7 (recombinant Avian)<sup>§</sup> | 48.5% (65.8%) [vs H7]* | 53.9% (58.7%) [vs N9]* |

| **Ferret Challenge Influenza A viruses (10xLD<sub>50</sub>) and Identity to Vaccine Viruses** |  |  |
| --- | --- | --- |
| Subtype (strain) | Percent nucleotide (and amino acid) identities |  |
|  | HA | NA |
| H1N1 (A/swine/Iowa/1931) | 77.6% (89.9%) | 85.5% (89.8%) |
| H2N7 (recombinant with 1957 Pandemic HA)<sup>§</sup> | 44.9% (42.5%) [vs H1]* | 53.9% (58.7) [vs N9]* |
| H3N2 (A/Port Chalmers/1973) | 83.8% (94.2%) | 43.5% (50.4) [vs N3]* |
| H10N7 (recombinant Avian)<sup>§</sup> | 48.5% (65.8%) [vs H7]* | 53.9% (58.7) [vs N9]* |

§Recombinant avian influenza A viruses made as 2:6 recombinants with above HA and NA gene segments with remaining 6 gene segments from A/green winged teal/Ohio/175/1986 (H2N1) [ref Qi].
\*Heterosubtypic HA and/or NA challenge virus homologies were compared to phylogenetically closest vaccine sequence as noted.

